# Metabolically-active obligate aerobes in anoxic (sulfidic) marine sediments

**DOI:** 10.1101/728287

**Authors:** Sabyasachi Bhattacharya, Chayan Roy, Subhrangshu Mandal, Moidu Jameela Rameez, Jagannath Sarkar, Svetlana Fernandes, Tarunendu Mapder, Masrure Alam, Rimi Roy, Nibendu Mondal, Prosenjit Pyne, Prabir Kumar Haldar, Aditya Peketi, Ranadhir Chakraborty, Aninda Mazumdar, Wriddhiman Ghosh

## Abstract

Metabolically-active obligate aerobes are unheard-of in tightly-anoxic environments. Present culture-independent and culture-dependent investigations revealed aerobic microbial communities along two, ~3-meter-long sediment-cores underlying the eastern Arabian Sea oxygen minimum zone, where high H_2_S disallows O_2_ influx from the water-column. While genes for aerobic respiration by *aa_3_*-/*cbb_3_*-type cytochrome-*c* oxidases and cytochrome-*bd* ubiquinol oxidase, and aerobic oxidation of methane/ammonia/alcohols/thiosulfate/sulfite/organosulfur-compounds, were present across the cores, so were live aerobic, sulfur-chemolithoautotrophs and chemoorganoheterotrophs. The 8820-years-old, highly–sulfidic, methane-containing sediment-sample from 275 cmbsf of 530 mbsl yielded many such obligately-aerobic bacterial-isolates that died upon anaerobic incubation with alternative electron-acceptors/fermentative-substrates. Several metatranscriptomic reads from this sediment-sample matched aerobic-respiration-/oxidase-reaction-/transcription-/translation-/DNA-replication-/membrane-transport-/cell-division-related genes of the obligately-aerobic isolates, thereby corroborating their active aerobic metabolic-status *in situ*. Metagenomic and metatranscriptomic detection of perchlorate-/chlorate-reduction genes, plus anaerobic growth of an obligately-aerobic *Halothiobacillus* isolate in the presence of perchlorate and perchlorate-reducing-consortia, suggested that cryptic O_2_ produced by perchlorate-respirers could be sustaining obligately-aerobes in this environment.

---

Geomicrobial dynamics of marine oxygen minimum zones (OMZs) have profound influence on oceanic biodiversity, productivity and fixed-nitrogen loss^1^. For the OMZs distributed across the global ocean, microbial community architectures of their anoxic sediments^2,3^ are less explored as compared to those of the hypoxic waters^4,5^. Whereas the water-columns of OMZs afford pM-nM O_2_, which is reportedly sufficient for successful maintenance of aerobic lifestyle^5–13^, their sediments have extremely shallow O_2_-penetration depth^14^ due to high flux of labile organic matter across the sea-bed leading to rapid consumption of O_2_ ^(15)^. As physicochemical considerations ruled out influx of O_2_ from the water column below a few centimeters of OMZ sea-floors^14^, potential ecology of aerobic microorganisms was never investigated in these anoxic sediments, despite the fact that plausible aerobic activities can have significant effects on the catabolic breakdown (remineralization) of the copious, labile organic matters that are sequestered in the sediments underlying hypoxic marine waters^15,16^.

Across the sediment-horizons of Arabian Sea, which harbors the thickest and one of the most intensely hypoxic marine OMZ^17^, maximum depth of O_2_ penetration reported is 1.6 centimeters below the sea floor (cmbsf)^14^. In a recent study based on eight ~3-m-long gravity-cores, collected on-board *RV* Sindhu Sankalp (SSK42) from across a water-depth transect covering the entire vertical-expanse of the eastern Arabian Sea OMZ (ASOMZ) (Supplementary Figure 1), we had revealed that anaerobic processes of the carbon-sulfur cycle heighten in the sulfidic sediments underlying the center of the OMZ’s vertical-expanse^2^. Here we explore the same SSK42-cores from the ASOMZ-center, namely SSK42/5 and SSK42/6 located at 580 and 530 meters below the sea-level (mbsl) respectively, for the potential ecology of aerobic bacteria. We used a “geochemistry - metagenomics – pure/mixed culture characterization – genomics - metatranscriptomics” approach to investigate microbiome structure/function in this sediment-horizon, where high concentrations of pore-water H_2_S^(2)^ preclude the presence/influx of free (gaseous/dissolved) O_2_, thereby making them ideal environments for exploring whether a functional ecology of aerobic bacteria is possible in a tightly anoxic environment.

## Results

### Geological context of the sample-site

Radiocarbon(^14^C)-based geological ages of the samples collected along SSK42/5 and 6 were found to range approximately between 1,000-12,000 and 4,000-10,600 yr BP respectively. Sedimentation has occurred in this region at rates ranging between 11-132 cm ky^−1^, without any apparent slumping or age reversal (Fig. 1a, e). Total organic carbon (TOC) content along SSK42/5 and 6 ranges between 1.2-4.6 and 0.5-3.7 wt % respectively^2^ (Fig. 1b, f). TOC content along either core does not correlate with sedimentation rate, thereby discounting down-depth dilution as a plausible reason for decrease in TOC content along SSK42/5 or 6; instead, the latter is attributable to temporal changes in TOC flux and/or organic matter degradation due to sulfate reduction^2^. None of the cores showed any visible sign of bioturbation. H_2_S, along SSK42/5, has a maxima of 427 μM at 105 cmbsf; in SS42/6, it increases with sediment-depth and reaches a maxima of 2010 μM at 250 cmbsf^(2)^ (see also Fig. 1c, g). Albeit technical limitations disallowed measurement of *in situ* O_2_ concentrations along SSK42/5 and 6, theoretical considerations indicated that there is no possibility of any free O_2_ remaining in, or diffusing into, the chemical milieu where H_2_S has already built-up, because the standing high-concentration of the strong reductant (H_2_S) would have already reacted with the strong electron acceptor (O_2_), either inorganically (following equation 1) or biochemically (following equations 2 and 3)^18^. Close to the sediment-surface of SSK42/5 and 6, concentration of dissolved O_2_ in the sea-water is approximately 2 µM, as measured by a CTD profiler^2^. This implied that a maximum of 2 µM O_2_ is available *in situ* for potential diffusion down the sediment pores, while the stoichiometries of the O_2_-H_2_S reactions given in equations 1–3 indicate that 1-4 µM *in situ* H_2_S would be sufficient to remove this O_2_ as it tries to diffuse down the sediment-pores. Notably, a build-up of 62 and 321 µM pore-water H_2_S was detected at 1 and 30 cmbsf of SSK42/5 and 6 respectively (only 0 and 15 cmbsf sediment-samples of SSK42/6 were found to be dissolved-sulfide-free), while deeper-down the two cores H_2_S-accumulation was even higher^2^. These H_2_S concentrations are high enough to scavenge not only the O_2_ diffusing into the sediments from the water-column, but also that which may be formed from potential abiotic/biotic processes and released directly into the chemical milieu. It can, therefore, be safely assumed that there is no dissolved O_2_ in the pore-waters beyond a few cmbsf, and a few tens of cmbsf, along SSK42/5 and 6 respectively.

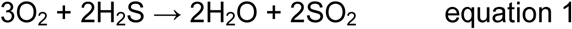

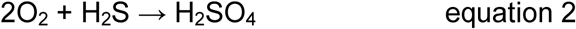

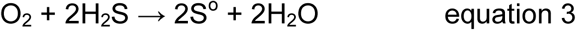

**Figure 1.**
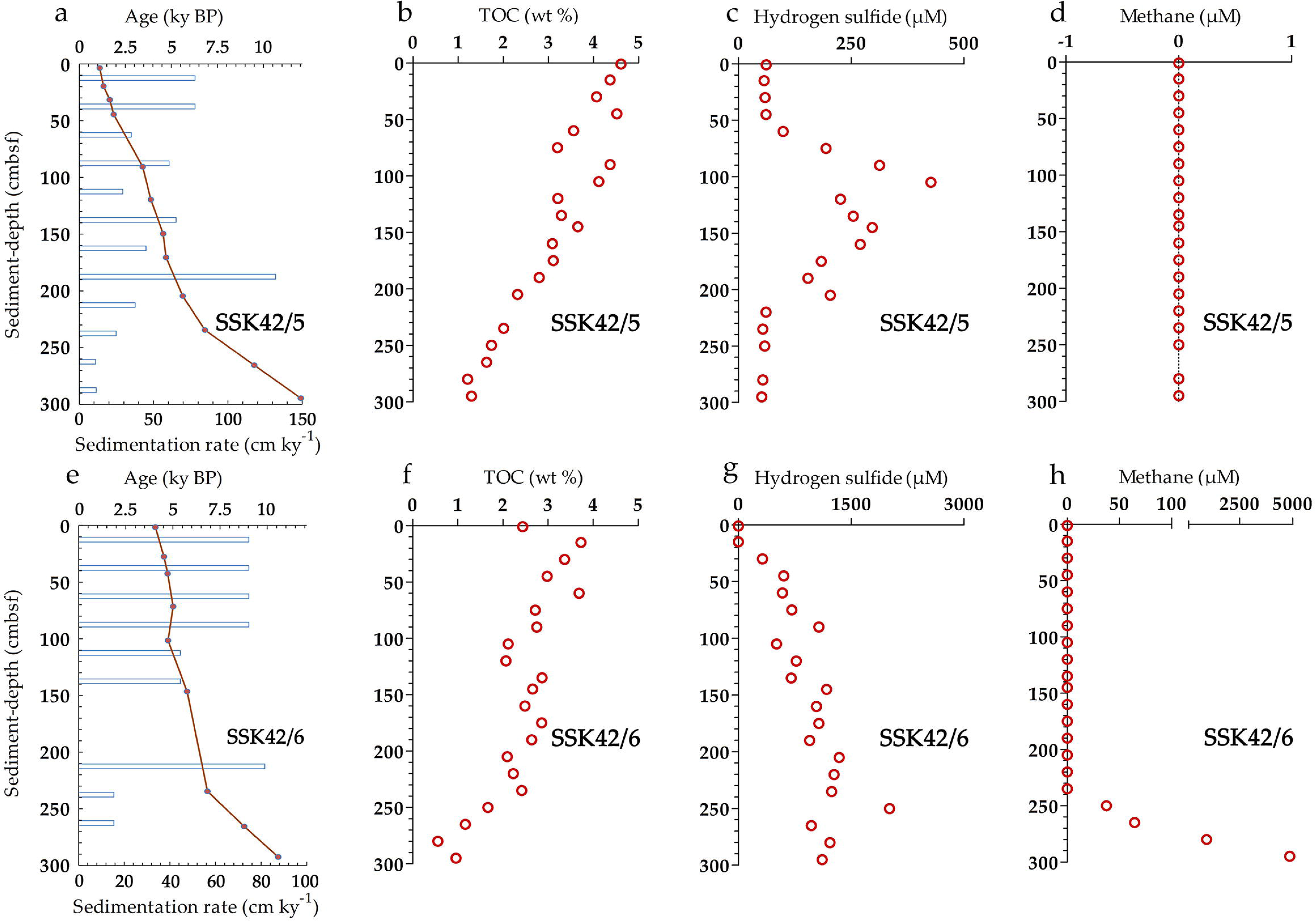
Age versus depth models and sedimentation rates, total organic carbon (TOC) contents (in wt %), hydrogen sulfide concentrations (in µM) and methane concentrations (in µM) along the cores (**a**-**d**) SSK42/5 and (**e**-**h**) SSK42/6. Sedimentation rates in both the cores were found to increase at around 6800 yr BP, and remain high towards the core-tops. Data for TOC, hydrogen sulfide and methane were taken from Fernandes et al. (2018)^2^.

### Genes governing O_2_-requiring metabolisms are abundant in the anoxic/sulfidic sediments of ASOMZ

Whole metagenome shotgun sequencing and analysis along the two cores revealed footprints of such metabolisms which cannot operate in the absence of O_2_. When the metagenomic sequence data obtained for each of the 25 distinct sediment-samples of SSK42/5 and 6 (Supplementary Tables 1, 2) were assembled and annotated individually, all the 25 contig-collections obtained were found to contain genes for aerobic respiration by *aa_3_*-type cytochrome *c* oxidase (*coxABCD*), *cbb_3_*-type cytochrome *c* oxidase (*ccoNOPQ*) and/or cytochrome-*bd* ubiquinol oxidase (*cydABX* and *appX*) (Supplementary Table 3). 19 out of the 25 contig-collections were found to contain genes for aerobic oxidation of methane/ammonia/alcohols [these included genes encoding particulate methane monooxygenase (*pmoABC*), soluble methane monooxygenase (*mmoXYZBCD*), methanol dehydrogenase (*mxaFI*) and/or alcohol oxidase (*mox*)] (Supplementary Table 4), whereas all 25 contained genes for the aerobic oxidation of thiosulfate (*soxC* for sulfane dehydrogenase), sulfite (SUOX for sulfite:acceptor oxidoreductase) and/or various organosulfur compounds (*dmoA* for dimethyl-sulfide monooxygenase, *ssuD* for alkanesulfonate monooxygenase, *sfnG* for dimethylsulfone monooxygenase and *mtoX* for methanethiol oxidase) (Supplementary Table 5).

In order to explore whether genes known to be involved in O_2_-dependent metabolisms are also present in other sulfidic/anoxic sediment-horizons of the sea, similar analyses as those described above were performed upon the H_2_S-containing sediment-samples of another 255-cm-long core, SSK42/9 (Supplementary Table 6). This comparator core was collected, during the same research cruise SSK42, from a shallower water-depth (31 mbsl) located outside the perennial ASOMZ (Supplementary Figure 1), ~128 km away from SSK42/5 and 6. When the metagenomic sequence data obtained from 10 discrete sediment-samples of SSK42/9 (Supplementary Table 6) were individually assembled and annotated, eight out of the 10 contig-collections were found to contain genes for aerobic respiration by the above mentioned low- and high-O_2_-affinity cytochrome oxidases (Supplementary Table 7). Four and five contig-collections from SSK42/9 also encompassed genes for aerobic oxidation of methane/ammonia/alcohols (Supplementary Table 8) and thiosulfate/sulfite/organosulfur-compounds (Supplementary Table 9) respectively. These data indicated that the phenomenon of aerobic metabolisms in sulfidic/anoxic sediments could be widespread in the marine realm.

### Enrichment, isolation and characterization of obligately aerobic bacterial strains from ASOMZ sediment

From the 275 cmbsf sediment-sample of SSK42/6 (age: 8820.3 yr BP), where ~1200 μM hydrogen sulfide and ~950 μM methane in the pore-waters^2^ (see also Fig. 1g, h) rule out the presence/influx of free O_2_ in the chemical milieu, aerobic microbial consortia could be successfully enriched in the following bacteriological media – (i) chemoorganoheterotrophic, Luria-Bertani (LB) broth, and Reasoner’s 2A (R2A) broth; (ii) chemolithoautotrophic, artificial sea water supplemented with thiosulfate (ASWT); and (iii) mixotrophic, modified basal and mineral salts-thiosulfate-yeast extract (MSTY) broth, and ASWT-yeast extract (ASWTY) broth. Simultaneously, high “most probable numbers” (MPN) of aerobic chemoorganoheterotrophs and aerobic sulfur-chemolithoautotrophs (determined in LB and ASWT respectively) were revealed along SSK42/5 and 6 (Supplementary Tables 10-13), thereby indicating that the aerobic bacteria of ASOMZ sediments, whether obligate or facultative, are alive *in situ*. 27 bacterial strains were isolated aerobically from the above mentioned enrichments in ASWT, MSTY and ASWTY. Taxonomically, the isolates formed nine species-level clusters, of which seven were classified under distinct genera while two clusters isolated separately in MSTY and ASWTY represented two distinct species of *Halomonas* (Table 1). One representative strain from each cluster was tested for anaerobic growth in its specific medium supplemented with the electron acceptors NaNO_3_, Fe_2_O_3_, MnO_2_, Na_2_SO_4_, (CH_3_)_2_SO and (CH_3_)_3_NO, which are all known to act as respiratory substrates in biogeochemically diverse marine microbiomes^19–21^. Subsequently, all the representative strains were also tested for anaerobic growth in their specific media supplemented with each of the above six electron acceptors separately. Furthermore, since many facultatively anaerobic or fermenting bacteria incapable of direct ferric iron reduction are known to channel their electrons towards iron reduction via external acceptors such as humic acids (this complex molecular mechanisms is called extracellular electron transfer)^22,23^, each of the current isolates were tested for anaerobic growth in their specific media supplemented with humic acids, or humic acids and Fe_2_O_3_ in combination. Aerobically incubated versions of all the above cultures exhibited cellular growth yields comparable with those obtained during aerobic growth in the same media-types minus any anaerobic respiratory substrate (Supplementary Table 14). This showed that the above mentioned electron acceptors or their used concentrations were non-toxic for the new isolates.

**Table 1.** The nine bacterial species isolated from 275 cmbsf of SSK42/6, and the anaerobic/fermentative growth properties of their representative strains.

|  | Bacteria isolated in ASWT |  | Bacteria isolated in ASWTY |  | Bacteria isolated in MSTY |  |  |  |  |
| --- | --- | --- | --- | --- | --- | --- | --- | --- | --- |
| Identification up to lowest taxonomic level possible | *Halothiobacillus* sp. | *Methylophaga* sp.* | *Sulfitobacter* sp. | *Halomonas* sp. | *Stenotrophomonas* sp. | *Halomonas* sp. | *Pusillimonas ginsengisoli* | *Pseudomonas bauzanensis* | *Cereibacter changlaensis* |
| Total number of strains isolated | 6 | 2 | 8 | 2 | 1 | 3 | 2 | 2 | 1 |
| Name of the representative strain | SB14A | SB9B<br>= MTCC 12599 | 15WGC<br>= MCC 3606 | 15WGF<br>= MCC 3301 | SBPC3 | SBBP1 | SBO3<br>= MTCC 12559 | SBBB<br>= MTCC 12600 | SBBC<br>= MTCC 12557 |
| 16S rRNA gene / whole genome sequence accession of the representative strain | LN999387 / SWAW01000000 | LN999390 / SSXS01000000 | LT607023 / SZNL01000000 | LT607031 / SSXT01000000 | LN999400 | LN999401 | LN999404 | LN999396 / SWAV01000000 | LN999397 / SWAU01000000 |
| **Anaerobic growth/survival in liquid cultures using electron acceptors other than O<sub>2</sub>** |  |  |  |  |  |  |  |  |  |
| Media used to check anaerobic growth<sup>§</sup> | ASWT | ASWM | ASWTY | ASWTY | MSTY | MSTY | MSTY | MSTY | MSTY |
| CFU<sup>#</sup> present mL<sup>-1</sup> culture at 0 h incubation | 6*10<sup>4</sup> | 6.1*10<sup>4</sup> | 2*10<sup>4</sup> | 3.8*10<sup>4</sup> | 7.8*10<sup>4</sup> | 4.5*10<sup>4</sup> | 1.9*10<sup>4</sup> | 5.1*10<sup>4</sup> | 2.6*10<sup>4</sup> |
| CFU<sup>#</sup> present mL<sup>-1</sup> culture after 10 days incubation | 2.8*10<sup>3</sup> | 0 | 0 | 0 | 1.6*10<sup>7</sup> | 9*10<sup>7</sup> | 2.3*10<sup>7</sup> | 0 | 0 |
| **Fermentative growth/survival in liquid cultures incubated under anaerobic condition** |  |  |  |  |  |  |  |  |  |
| Media used to check fermentative growth | ASWT | ASWMF | ASWTYP | ASWTYP | NA | NA | NA | MSTYP | MSTYP |
| CFU<sup>#</sup> present mL<sup>-1</sup> culture at 0 h incubation | 7.9*10<sup>4</sup> | 7.8*10<sup>4</sup> | 2.2*10<sup>4</sup> | 3.5*10<sup>4</sup> | NA | NA | NA | 6*10<sup>4</sup> | 2.5*10<sup>4</sup> |
| CFU<sup>#</sup> present mL<sup>-1</sup> culture after 5 days incubation | NR | 2.8*10<sup>4</sup> | 0 | 0 | NA | NA | NA | 1.6*10<sup>3</sup> | 0 |
| CFU<sup>#</sup> present mL<sup>-1</sup> culture after 15 days incubation | 2.8*10<sup>3</sup> | 0 | NA | NA | NA | NA | NA | NR | NA |
| CFU<sup>#</sup> present mL<sup>-1</sup> culture after 45 days incubation | NR | NA | NA | NA | NA | NA | NA | 0 | NA |
| CFU<sup>#</sup> present mL<sup>-1</sup> culture after 60 days incubation | 3.7*10<sup>2</sup> | NA | NA | NA | NA | NA | NA | NA | NA |
| CFU<sup>#</sup> present mL<sup>-1</sup> culture after 80 days incubation | 4.4*10<sup>1</sup> | NA | NA | NA | NA | NA | NA | NA | NA |
| CFU<sup>#</sup> present mL<sup>-1</sup> culture after 100 days incubation | 0 | NA | NA | NA | NA | NA | NA | NA | NA |

\* While all isolates were maintained, and tested for anaerobic growth, in their respective isolation-media, *Methylophaga* was maintained/tested in ASW-methanol (ASWM).
MSTYP = MSTY medium supplemented with pyruvate; ASWTYP = ASWTY medium supplemented with pyruvate; ASWMF = ASWM medium supplemented with fructose; full-forms of all the other abbreviated media names are available in the text. NA = Not applicable; NR = Not recorded.
§ All media-types used to check anaerobic growth/survival were supplemented with MnO<sub>2</sub>, Na<sub>2</sub>SO<sub>4</sub>, NaNO<sub>3</sub>, Fe<sub>2</sub>O<sub>3</sub>, (CH<sub>3</sub>)<sub>2</sub>SO and (CH<sub>3</sub>)<sub>3</sub>NO.

After 10 days of anaerobic incubation in their specific media supplemented with NaNO_3_, Fe_2_O_3_, MnO_2_, Na_2_SO_4_, (CH_3_)_2_SO and (CH_3_)_3_NO,the three representative strains MSTY-isolated*-Halomonas* SBBP1, *Pusillimonas ginsengisoli* MTCC 12559 and *Stenotrophomonas* sp. SBPC3 exhibited growth (Table 1). These three strains could also grow anaerobically in their respective media when NaNO_3_, but not the other five compounds, were provided individually as the sole electron acceptor - their cellular growth yields in the presence of NaNO_3_ were identical to those obtained in the presence of the six-electron-acceptor mixture (Supplementary Table 15). In contrast to the above three strains, after 10 days of anaerobic incubation in their specific media containing the six-electron-acceptor mixture, the representative *Halothiobacillus* strain SB14A retained only 4.6% of the initial cell count whereas the representative strains of the *Cereibacter*, ASWTY-isolated-*Halomonas*, *Methylophaga*, *Pseudomonas* and *Sulfitobacter* clusters had no viable cells left (Table 1). Use of the six individual electron acceptors separately in six different culture sets did not also help any of these strains to grow anaerobically (Supplementary Table 15). These results were consistent with the fact that no genetic system for respiration using nitrate, ferric, manganic or sulfate ion, or dimethyl sulfoxide or trimethylamine N-oxide, were detected when the *de novo* sequenced and annotated genomes of these representative strains, namely *Halothiobacillus* sp. SB14A, *Cereibacter changlaensis* MTCC 12557, ASWTY-isolated *Halomonas* sp. MCC 3301, *Methylophaga* sp. MTCC 12599, *Pseudomonas bauzanensis* MTCC 12600 and *Sulfitobacter* sp. MCC 3606 were scrutinized manually as well as using the *KEGG (Kyoto Encyclopedia of Genes and Genomes) Mapper - Reconstruct Pathway* tool hosted by Kanehisa Laboratories at https://www.genome.jp/kegg/tool/map_pathway.html.

After 10 days of anaerobic incubation in their specific media supplemented with humic acids, or humic acids and Fe_2_O_3_ in combination, neither the three strains that were characterized above as facultative anaerobes, nor those six which were found to be obligate aerobes, had any viable cell left in the cultures (Supplementary Table 15). Consistent with these findings, none of the key genes related to extracellular electron transfer (e.g., those encoding surface proteins such as Cyc2, a monoheme cytochrome *c*, and multiheme *c*-type cytochromes; porin-cytochrome *c* protein complexes in which multiheme *c*-type cytochromes are secreted to the periplasm and embedded into a porin on the outer membrane to form the extracellular electron transfer conduit^23^) were detected upon manual as well as *KEGG Mapper - Reconstruct Pathway*-based scrutiny of the annotated genomes of the six obligately aerobic isolates.

When the *Cereibacter*, ASWTY-isolated-*Halomonas*, *Pseudomonas* and *Sulfitobacter* strains were tested for fermentative growth/survival in their specified media supplemented with pyruvate, none, except the *Pseudomonas* strain, had any viable cell left after 5 days of anaerobic incubation. The *Pseudomonas* strain too had no viable cell left after 45 days. Fermentative survival of the *Methylophaga* strain was tested in ASW-methanol-fructose^24^, and that of the *Halothiobacillus* strain in ASWT [known strains of *Halothiobacillus* cannot utilize organic carbon, so this was to test survival via fermentation of stored polyglucose^25^] - the two strains had no viable cells left after 15 and 100 days respectively (Table 1). Corroborative to these data, genomes of the six obligately aerobic strains encompssed no gene encoding for the enzymes (such as lactate dehydrogenase, pyruvate:ferredoxin oxidoreductase, NADH: ferredoxin oxidoreductase, ferredoxin dehydrogenase, phosphotranscetylase, acetate kinase, acetaldehyde dehydrogenase, and acetyly-coA acetyl transferase) that are required for transformations of pyruvate to glycolytic fermentations.

### *De novo* sequencing and analysis of the genomes of the obligately aerobic isolates

Whole genome shotgun sequencing, assembly and annotation were carried out for the six obligately aerobic isolates *Cereibacter changlaensis* MTCC 12557, *Halomonas* sp. MCC 3301, *Halothiobacillus* sp. SB14A, *Methylophaga* sp. MTCC 12599, *Pseudomonas bauzanensis* MTCC 12600 and *Sulfitobacter* sp. MCC 3606 (GenBank accession numbers are given in Table 1; general features of the assembled genomes given in Supplementary Table 16). All the six genomes were found to contain several oxidase-encoding genes, including those for *cbb_3_*-type cytochrome *c* oxidase. All except *Methylophaga* sp. MTCC 12599 contained *aa_3_*-type cytochrome *c* oxidase genes, while all except *Halothiobacillus* sp. SB14A had genes for cytochrome-*bd* ubiquinol oxidase (see Supplementary Tables 17-22 in the alphabetical order of the six isolates).

Significant prevalence of all the six obligately-aerobic strains was evident across the OMZ sediment-horizon when substantial percentages of the metagenomic reads available for the individual sample-sites of SSK42/5 and 6 mapped onto each of the six newly-sequenced genomes (Fig. 2; Supplementary Table 16). In SSK42/5 and 6, approximately 0.01-0.3% and 0.04-19.05% metagenomic reads from the individual sample-sites mapped onto the six different genomes respectively; the only metagenome-genome pair to have no matching sequence was “45 cmbsf of SSK42/6 and *Cereibacter changlaensis* MTCC 12557”. Prevalence of reads matching the six new isolates was relatively higher for the metagenomes of SSK42/6 (Fig. 2), and then within this core, highest for 275 cmbsf, the sample-site from where the strains were isolated (Supplementary Table 16). Furthermore, in this context, it is noteworthy that much higher proportions (0.6-1.0%) of metagenomic reads from the 0-115 cmbsf sample-sites of the comparator, H_2_S-containing sediment-core SSK42/9 mapped onto each of the six assembled genomes, as compared to the corresponding values for entire SSK42/5. Below 115 cmbsf of SSK42/9, however, metagenomic reads matching these genomes were almost absent (0.01-0.05% metagenomic reads from the 125-255 cmbsf sample-sites of SSK42/9 mapped onto the six individual genomes; see Fig. 2; Supplementary Table 16).

**Figure 2.**
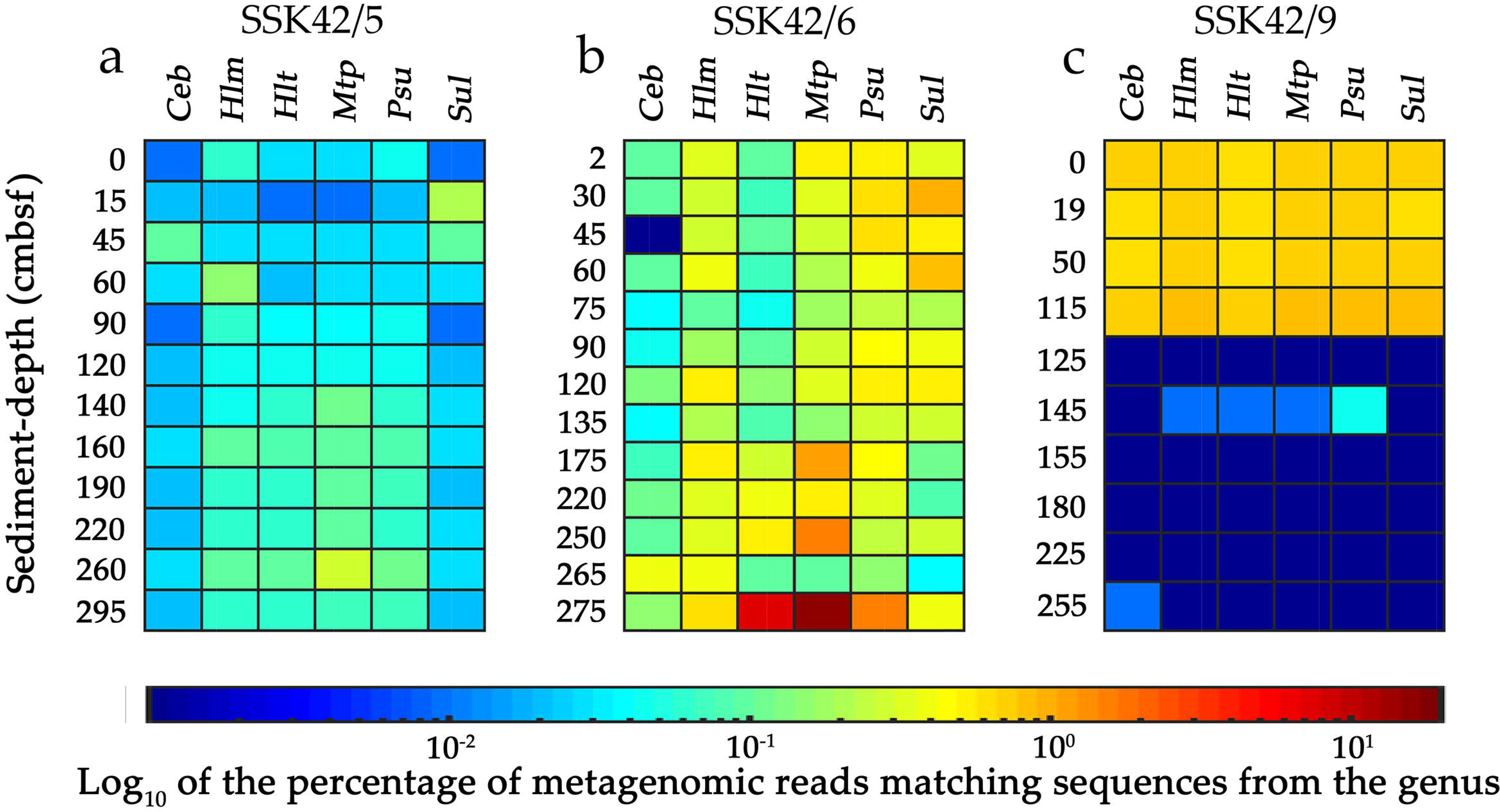
Heat map comparing the percentages of metagenomic reads from individual sediment-samples of (**a**) SSK42/5, (**b**) SSK42/6 and (**c**) SSK42/9 that matched with genomic sequences of the six obligately aerobic bacterial isolates: (*Ceb*) *Cereibacter changlaensis* MTCC 12557, (*Hlm*) *Halomonas* sp. MCC 3301, (*Hlt*) *Halothiobacillus* sp. SB14A, (*Mtp*) *Methylophaga* sp. MTCC 12599, (*Psu*) *Pseudomonas bauzanensis* MTCC 12600, and (*Sul*) *Sulfitobacter* sp. MCC 3606. For each percentage value, its Log_10_ has been plotted in the z axis of the heat map, so as to resolve the wide span of the data. Percentage level of matched reads for individual pairs of metagenomic-genomic datasets ranged between 0 and 19.05, with intermediate values in the order of 10^−2^ to 10^1^. Since 0 is less than any minimum value possible in the log scale, it is noteworthy that the deepest blue squares of panels B and C, which apparently match the order of 10^−3^ in the color scale, actually correspond to the real values 0 (see Supplementary Table 16).

### Correspondence between genomic sequences of the obligately aerobic isolates and metatranscriptomic sequences from their habitat reflects *in situ* aerobic activity

The six bacterial isolates which not only failed to grow but also died within a few days/months of anaerobic incubation (despite being provided with alternative electron acceptors or fermentative substrates; see Table 1), could not have remained alive for thousands of years in their natural habitat without respiring O_2_, whatever may be its source and concentration *in situ*. There may well be a debate regarding which of the three metabolic states broadly referred to as living (namely dormant; active but not growing; and active and growing)^26^ did these strains exist in, for thousands of years in the ASOMZ sediment. In any case, it is for certain that none of the three states of life could have been achieved without some level of metabolic activity (to support basic cell maintenance, activity and/or growth)^27–29^, and any metabolic activity in an obligate aerobe ought to eventually involve aerobic respiration. However, since *in situ* aerobic metabolism is counterintuitive to the physicochemical conditions prevailing in the habitat (Fig. 1), we considered it imperative to test whether the native metatranscriptome of the sediment-sample from where the obligately aerobic strains were isolated contained transcripts corresponding to the aerobic metabolism-related genes of those strains. Furthermore, previous comparative analysis of total-mRNA sequence data from ecologically diverse microbial communities had identified that transcripts corresponding to core metabolic genes such as those associated with carbohydrate and energy metabolism, amino acid metabolism, nucleotide metabolism, protein translocation and cell division, are present in all metatranscriptomes^30^. Presence of such mRNAs were not only considered to be the quality/coverage benchmarks of the metatranscriptomic datasets, but also regarded as markers of *in situ* metabolic activity of the source bacteria^30^. We, therefore, additionally tested whether the native metatranscriptome from 275 cmbsf of SSK42/6 contained mRNA sequences corresponding to these metabolic-activity markers of the obligately aerobic isolates.

Whole metatranscriptome of the 275 cmbsf sediment-sample of SSK42/6 was sequenced, and the paired end mRNA reads obtained were mapped onto the manually-curated aerobic-metabolism-related gene-catalogs of the six obligately aerobic strains (Supplementary Tables 17-22). Maximum number of metatranscriptomic read-pairs mapped onto the aerobic-metabolism gene-catalog of *Halothiobacillus* sp. SB14A, with a total 7492 read-pairs (i.e., 0.03% of the total 26,496,769 read-pairs examined) concordantly matching genes for *aa_3_*-type cytochrome *c* oxidase, *cbb_3_*-type cytochrome oxidase, and various other oxidase enzymes that, as a group, are all characterized by the catalysis of redox reactions involving O_2_ as the electron acceptor (Supplementary Table 19). For *C*. *changlaensis* MTCC 12557 (Supplementary Table 17), *Halomonas* sp. MCC 3301 (Supplementary Tables 18), *P*. *bauzanensis* MTCC 12600 (Supplementary Table 21) and *Sulfitobacter* sp. MCC 3606 (Supplementary Table 22), metatranscriptomic read-pairs matched concordantly with their genes for *aa_3_*-type cytochrome *c* oxidase, *cbb_3_*-type cytochrome oxidase, cytochrome-*bd* ubiquinol oxidase, as well as various other oxidases - total number of read-pairs mapping onto the aerobic-metabolism-related gene-catalogs of these four strains were 665, 2327, 4845 and 188 respectively (these numbers were 0.003, 0.009, 0.02 and 0.0007 percent of the total 26,496,769 read-pairs tested). For *Methylophaga* sp. MTCC 12599, only 13 metatranscriptomic read-pairs (which was only 0.00005% of the total read-pairs tested) matched concordantly with its gene for cytochrome-*bd* ubiquinol oxidase subunit II (Supplementary Table 20).

Whether the six obligately aerobic strains were metabolically active *in situ* was further corroborated by mapping the metatranscriptomic reads obtained from the 275 cmbsf sediment-sample of SSK42/6 onto their gene-catalogs pertaining to the metabolic categories (**i**) **<u>G</u>enetic Information Processing** (*Transcription*: RNA polymerase, *Translation*: Ribosome and Aminoacyl-tRNA biosynthesis, *Replication and repair*: DNA replication); (**ii**) **<u>E</u>nvironmental Information Processing** (*Membrane transport*: ABC transporters, Phosphotransferase system and Bacterial secretion system) and (**iii**) **<u>C</u>ell Growth and Division** (Cell cycle), abbreviated here onwards as **<u>GEC</u>** gene catalogs. Of the six obligately aerobic isolates, *Halothiobacillus* sp. SB14A had the maximum number (221749) of metatranscriptomic read-pairs (i.e., 0.84% of the total 26,496,769 read-pairs tested) matching concordantly with 181 genes of its 183-gene strong GEC catalog (Supplementary Table 23). For *C*. *changlaensis* MTCC 12557 (Supplementary Table 24), *Halomonas* sp. MCC 3301 (Supplementary Table 25), *P*. *bauzanensis* MTCC 12600 (Supplementary Table 26) and *Sulfitobacter* sp. MCC 3606 (Supplementary Table 27), 22415, 247934, 291711 and 33863 metatranscriptomic read-pairs (i.e., 0.09, 0.94, 1.10 and 0.13% of the total 26,496,769 read-pairs examined) matched concordantly with 245, 132, 147 and 133 genes of their 397-, 273-, 252- and 310-gene strong GEC catalogs. For *Methylophaga* sp. MTCC 12599, only 7327 metatranscriptomic read-pairs (which was only 0.03% of the total read-pairs examined) matched concordantly with 34 genes of its 193-gene strong GEC catalog (Supplementary Table 28).

### Perchlorate-/chlorate-respiring bacteria as a potent source of O_2_ for aerobic bacteria in ASOMZ sediments

In view of the absence of free O_2_ in the sulfidic chemical milieu of SSK42/5 and 6, it was imperative to explore the potential source(s) of cryptic O_2_ that may be facilitating aerobic life *in situ*. Perchlorate/chlorate disproportionation^31^ is a microbial metabolic process that can potentially serve this purpose. Perchlorate (ClO_4_^−^) and chlorate (ClO_3_^−^) respiring bacteria disproportionate the highly oxidized states of chlorine to chloride (Cl^−^) and O_2_ by the consecutive actions of (per)chlorate reductase (that catalyzes the reduction of perchlorate to chlorite via chlorate) and chlorite dismutase (that converts chlorite to chloride and oxygen)^31^. Remarkably, several genes encoding (per)chlorate reductase and/or chlorite dismutase were identified within the metagenome assemblies carried out for all the individual sediment-samples of SSK42/5 and 6, except 220 and 250 cmbsf of SSK42/6 (Supplementary Table 29). We, therefore, hypothesized that potentially-close association with these oxygenic anaerobes could be one of the means by which aerobic lifestyle is sustained in the ASOMZ sediments. Furthermore, when the metagenomic sequence data obtained from the 10 discrete sediment-samples of SSK42/9 were individually assembled and annotated, 8 out of 10 contig-collections obtained were also found to contain (per)chlorate reductase genes (Supplementary Table 30).

Guided by the above hypothesis, perchlorate-respiring bacterial consortia (PRBC) could be successfully enriched in perchlorate-supplemented, basal bicarbonate-buffered (PBBB) medium by anaerobically incubating (at 15°C for 45 days) different sediment-samples of SSK42/5 and 6. Following another round of 7 day sub-culturing, enrichments corresponding to 120 and 260 cmbsf of SSK42/5, and 135 and 275 cmbsf of SSK42/6, produced maximum O_2_ and/or biomass; so these PRBC were selected for individual anaerobic co-culture experiments with the obligately aerobic isolate *Halothiobacillus* SB14A in a PBBB-ASWT consensus medium containing energy/electron/carbon sources suitable for both biotypes, but only ClO_4_^−^, and no O_2_, as electron acceptor. After 60 days of anaerobic incubation in the consensus medium, SB14A, when accompanied by a PRBC, exhibited 1.1-22 fold increases in viable-cell count (in the different co-culture sets, increases in viable-cell count of *Halothiobacillus* SB14A varied because the different O_2_-providing, undefined-consortia that were used contained differential diversities and abundances of perchlorate-respirers; this caused O_2_-availability, and therefore *Halothiobacillus* growth, to vary). In contrast, when SB14A was incubated anaerobically in the same medium, but in the absence of any PRBC, the culture lost 99.6% of the initial cell count (Supplementary Table 31). Furthermore, after 60 days of anaerobic incubation in a perchlorate-less variant of the consensus medium, SB14A cultures, whether accompanied by a PRBC or not, lost 99.6-99.7% of the initial cell counts (Supplementary Table 31). These results corroborated the feasibility of obligate aerobes surviving in anoxic habitats by means of potentially close associations with oxygenic, perchlorate-respiring bacteria.

No ClO ^−^ was detected in the pore-water of any sediment-depth explored along SSK42/5, 6 or 9 using ion chromatographic method that apparently had a detection limit of 10 µM (1 ppm). However, potentially functionality of perchlorate respiration in these sediments was suggested by the detection of (per)chlorate reductase homologs in the gene-catalog obtained via assembly and annotation of the metatranscriptomic sequence dataset obtained from 275 cmbsf of SSK42/6 (Supplementary Table 32). Presence of no chlorite dismutase homolog in the assembled-metatranscriptome-derived gene-catalog, or for that matter presence of relatively less number of homologs for chlorite dismutase than (per)chlorate reductase in the assembled-metagenome-derived gene-catalogs, could be reflective of the *in situ* prevalence of novel molecular diversities of chlorite dismutase genes which did not get detected in automated annotation procedures. This supposition is consistent with the fact that *de novo* sequencing and automated annotation of the genomes of perchlorate-respiring bacterial strains isolated subsequently from 275 cmbsf of SSK42/6 also revealed no chlorite dismutase homolog (Unpublished data of this research group).

## Discussion

In contemporary biogeochemical explorations of marine OMZ waters, <3 nM dissolved O_2_ has been regarded as anoxic due to the limitations of field-applicable O_2_-detection technologies^4,7,13^. It is, however amply illustrated in the literature that the low levels of dissolved O_2_ present in OMZ waters are not only supportive of *in situ* aerobic metabolisms^5,8,10,12,13^ but also reasonably within the known frontiers of minimum-O_2_ necessary for aerobic microbial life^9,11^. By contrast, the H_2_S-rich sediments underlying the center of the vertical expanse of the eastern ASOMZ are far more tightly anoxic. Revelation of live and metabolically active aerobic microorganisms in such an environment, therefore, opens up a new frontier in understanding Life’s opportunities and adaptations in the extremes.

Results obtained from the metatranscriptomic read mapping experiments showed that the obligately aerobic strains, in their native habitat, were not only metabolically active and respiring with O_2_, but were also growing/replicating in all likelihood. Inferring metabolic activity from transcript profiles is justified not only since mRNAs effect all metabolic functions and fluxes through their translation into enzymes and transporters, but also because, in the light of evolutionary optimality^32^, it is axiomatic that if a cell undertakes mRNA production it does so only to effect the function in which the coded protein is involved. As for environmental metatranscriptomes, instantaneous inventories of mRNA pools are considered to be highly informative about ecologically-relevant processes (community functions) going on *in situ*^33^, while annotation of total environmental mRNAs with reference to specific genome sequences are adept to revealing the active biochemical functions of particular microorganisms within a community^30^. mRNAs in eukaryotic and bacterial cells have half-lives in the order of hours to days, and only a few minutes to hours, respectively^34,35^. In bacteria, mRNA half-lives average between 1 and 8□min, for both laboratory-grown and environmental cells, independent of their growth rates^33,35^; for some extremophilic archaea, transcript stabilities are a bit higher and range between 5 and >18 min^36^. For marine bacterial isolates, half-life time of mRNA transcripts have been found to range between 1 and 46 min, while for coastal metatranscriptomes it ranges from 9 to 400 min^37^. Notably, mRNAs do not form secondary structures as tRNAs and rRNAs do, and therefore they are far more unstable than the latter types of RNA. This may the reason why rRNAs from some eukaryotic organisms have been found to remain stable over geological time periods, in cellular as well as extracellular milieus, across marine subsurface provinces^38^, whereas there is not a single report of long-term preservation of any prokaryotic mRNA in geological samples. Bacteria in various coastal environments, freshwaters, soils, and sediments have an average mRNA content of only 200-300 molecules per cell, and each of these last for only a few minutes before being degraded^33^; this allows the metabolic state of a cell to adjust to changing environmental conditions via alteration of its expressing gene-set^36^. Consequently, in an environmental sample, it is impossible to obtain mRNA transcripts of a bacterium (especially those corresponding to core metabolic activities such as transcription, translation, DNA replication, membrane transport and cell cycle) in case of its *in situ* dormancy or cell-system collapse (death). As for the 15-25°C sediments of SSK42/5 and 6, which are characterized by high *in situ* organic carbon content and hectic carbon-sulfur cycling^2^, we hypothesize, based on all the above considerations, that longevity of bacterial mRNA, in case of dormancy or cell death, could be in the order of few minutes. This supposition is corroborated by the reported identification of very low amount of mRNA in dormant laboratory cultures of *Mycobacterium tuberculosis*^39^, and revelation of strong positive correlation between the abundance of bacterial mRNAs involved in cell division and *in situ* concentration of bacterial cells cell, during the metatranscriptomic exploration of anaerobic Peru Margin sediments^40^. Notably, metatranscriptome analysis has also been used to delineate active prokaryotic community functions in O_2_-depleted sediments of Baltic Sea that are biogeochemically comparable with the ASOMZ sediments^41^.

Based on the data obtained in this study, the role of perchlorate-/chlorate-respiration as a cryptic source(s) of O_2_ for aerobic life in the anoxic sediments of ASOMZ is at best hypothetical. The bases of origin, concentration-levels, distribution, and bioavailability of ClO_4_^−^ (which is thus far appreciated mainly as a chemical of anthropogenic origin^42^) in the marine realm is completely unknown, and requires substantiation by new studies of biogeochemistry. Whilst there is every possibility of other biotic/abiotic contrivances being there in the ASOMZ sediment ecosystem to meet the O_2_ requirement of strictly aerobic life, and/or to take care of the electron load of the obligate aerobes living *in situ*, in the current context, it is noteworthy that a number of previous studies have reported the isolation of perchlorate-/chlorate-respiring bacterial strains (endowed with all genetic components necessary for perchlorate reduction) from marine sediment-samples that, like SSK42/5 and 6, also did not contain detectable perchlorate^43–45^. Furthermore, from a purely microbiological point of view, close symbiotic association of perchlorate-reducers with aerobic microorganisms (that can scavenge O_2_ for them) makes ecophysiological sense in the context of the anoxic ASOMZ sediments. Perchlorate-reducers respire O_2_ (and sometimes nitrate) in preference over perchlorate, and for some of them 2-6 mg L^−1^ O_2_ inhibits perchlorate reduction irreversibly^31^. However, in redox-stratified unbioturbated sediments^46^ as those of SSK42/5 and 6^(2)^, O_2_ is consumed within the top few millimeters, and nitrate penetrates slightly deeper than that; consequently, perchlorate-respiration, and therefore a low-O_2_ micro-environment, apparently becomes mandatory for the *in situ* survival of these bacteria.

Whatever may be the source(s) of cryptic O_2_ in the sulfidic/anoxic sediments underlying the eastern Arabian Sea OMZ, metabolically active obligate aerobes in such territories (Fig. 2) hold potent implications for the *in situ* remineralization of organic matter and oxidation of sulfur, iron and nitrogen species. OMZs and similar hypoxic marine water bodies are characterized by high flux of labile organic matter across the sea-bed^15,16^, and bulk of this is likely to be complex in nature (i.e. comprising large number of carbon atoms) because low O_2_ in the water columns^4,7,13^, and zero O_2_ below a few centimeters from the sea-floor^14^, disallows large-scale aerobic degradation. For the sediments of SSK42/5 and 6 too, TOC content is copious, and anaerobic processes of the carbon-sulfur cycle (that are fuelled by simple carbon compounds such as acetate, lactate, formate, methanol, etc.) are highly active^2^. So it had thus far been a biogeochemical intrigue as to how the high demand for simple carbon compounds is satisfied in this kind of sedimentary ecosystems which are impinged by acutely hypoxic waters. Irrespective of whether the simple carbon compounds produced from the low levels of aerobic catabolism in the hypoxic waters/sediment-surfaces (together with those potentially formed within the sediment package from plausible anaerobic hydrolysis and fermentation) are enough for this purpose, the present discovery of metabolically active crypto-aerobic microflora within this anoxic sediment-horizon clearly adds a whole new dimension to the scope and extent of *in situ* remineralization of sequestered organic carbon. By design, implications of this phenomenon are most likely to extend to carbon, sulfur, nitrogen and metals cycling in the benthic realm at large and plausibly also in other anoxic territories of the Earth’s biosphere. New lines of concerted investigation of the ecology of crypto-aerobic microbial communities (via omics-as well as culture-based approaches) and their *in situ* geological interactions, manifestations and records are needed to gain comprehensive insights into the global biogeochemical significance of aerobic microbial life amidst anoxia.

## Methods

### Sample collection and storage

The gravity-cores SSK42/5, 6 and 9 were collected from 580 (16°49.88’ N, 71°58.55’ E), 530 (16°50.03’ N, 71°59.50’ E) and 31 mbsl (16°13.56’ N, 73°19.16’ E) respectively, off the west coast of India, and sampled under constant shower of high-pure N_2_, as described previously^2^. Immediately after cutting and removing the longitudinal halves of the PVC core-liners, top one cm of the exposed surfaces were scrapped-off to avoid contaminations from the core-liner’s inner surface and/or sea water; sampling was then carried out from the interiors. For every sediment-depth explored, two sample-fractions designated for duplicate total community DNA/RNA isolation were collected and put into −20ºC freezers, while one sample-fraction each for culture-based microbiology and chemistry were kept at 4ºC. Before refrigeration, sample bottles were flushed with nitrogen gas, screw-capped, and sealed with Parafilm (Bemis Company Inc., USA). During transfer to the institute laboratories, and also over long-term sample-preservation, the above mentioned temperatures were maintained consistently. During investigation, sample-bottles meant for chemistry or culture-independent microbiology were always opened aseptically under constant nitrogen flow; sample-bottles meant for culture-based microbiology were opened inside a Whitley H35 Hypoxystation (Don Whitley Scientific, UK) preset at 75% humidity and 0% partial pressure of O_2_ using the gas mixture N_2_:H_2_:CO_2_ = 80:10:10 (v/v/v). After every use, sample-bottles were screw-capped and sealed with fresh Parafilm.

### Age of the samples and sedimentation rate

Radiocarbon (^14^C) dates were determined on bulk planktonic foraminifera. An assemblage of ~200 specimens was cleaned ultrasonically in sodium hexametaphosphate solution and H_2_O_2_ (30%) in order to remove any adhered clay particles and organic carbon respectively. After the specimens were rinsed with distilled water, Foraminifera tests were further broken to remove fragments bearing pyrite, quartz and authigenic carbonates as much as possible. ^14^C dates were determined using accelerator mass spectrometry (AMS) at the National Ocean Sciences AMS Facility of Woods Hole Oceanographic Institution, Woods Hole, MA, USA. Carbon dioxide generated from the dissolution of foraminifera shells in H_3_PO_4_ was reacted with catalyst to form graphite, which was then pressed into targets and analyzed on the accelerator along with standards and process blanks. NBS Oxalic Acid I (NIST-SRM-4990) and Oxalic Acid II (NIST-SRM-4990C) were used as the primary standards during ^14^C measurements^47^. Radiocarbon ages were converted to mean calendar ages using the calibration curve of Fairbanks et al. (2005)^48^. Ages were expressed as radiocarbon years before 1950 AD, or years before present (yr BP), using the Libby half-life of 5568 years. 400 years were used as reservoir age for calibration^49^.

### Extraction of total DNA/RNA from sediment-samples/microbial cultures

Total community DNA was extracted from native sediment-samples using PowerSoil DNA Isolation Kit (MoBio, USA) as per the manufacturer’s protocol. Microgram-levels of DNA were obtained from each preparatory reaction that started with 0.5 g sediment-sample. Genomic DNA of the pure culture isolates was extracted using HiPurA Bacterial Genomic DNA Purification Kit (Himedia, India), following manufacturer’s instructions. Total community RNA was extracted from the 275 cmbsf sediment-sample of SSK42/6 using the RNA PowerSoil Total RNA Isolation Kit (MoBio), as per manufacturer’s protocol. Nanogram-level total RNA was obtained after pooling up the products of 15 individual preparatory reactions, each carried out using 2 g sediment-sample.

### Sequencing of metagenomes/genomes/metatranscriptomes

The duplicate metagenomes isolated for each sediment-depth explored along SSK42/5, 6 and 9 were shotgun sequenced individually on an Ion Proton sequencing platform (Thermo Fisher Scientific, USA) using 200 nucleotide read-chemistry, as described previously^50^ (details given in Supplementary Methods; complete lists of sedimentary communities investigated along SSK42/5, 6 and 9 are given in Supplementary Tables 1, 2 and 6 respectively). Whole genomes of the pure culture isolates were shotgun sequenced on an Ion S5 platform (Thermo Fisher Scientific) using 400 nucleotide read-chemistry (details given in Supplementary Methods). The total community RNA extracted from the 275 cmbsf sediment-sample of SSK42/6 was sequenced on a HiSeq4000 platform (Illumina Inc., USA) using paired end, 2 × 150 nucleotide, sequencing by synthesis read-chemistry with dual indexing workflows. Details of sequencing-library preparation are given in Supplementary Methods. Depletion of rRNAs was carried out using Ribo-Zero Gold (Illumina), which is an integral part of the TruSeq Stranded mRNA and Total RNA kit (Illumina) used for preparing the library. Very little rRNA escaped the Ribo-Zero treatment where, according to the kit manual, biotinylated probes bind rRNA species selectively, and magnetic beads capture and pull-down the probe-rRNA hybrids, leaving the desired rRNA-depleted RNA pool in solution. In order to extract and eliminate whatever rRNA reads were there in the raw metatranscriptomic sequence dataset, the 26,579,343 read-pairs available in all were mapped onto SILVA large subunit as well as small subunit rRNA gene sequence database^51^, using the short read aligner Bowtie2 v.2.3.4.3^(52)^ in default local (sensitive) alignment mode; this identified only ~0.3% metatranscriptomic reads as ascribable to rRNAs, thereby leaving 26,496,769 read-pairs in the rRNA-sequence-free final dataset used for downstream analyses (notably ~20 million reads has been proven reasonably sufficient for enzyme discovery in any microbiome in general)^30^.

### *De novo* assembly and annotation of metagenomes/genomes/metatranscriptome

All sequence datasets were quality-filtered with Phred score cut-off 20 using Fastx_toolkit 0.0.13.2 (http://hannonlab.cshl.edu/fastx_toolkit/download.html). High quality reads from the duplicate metagenomic sequence datasets available for each sediment-community were co-assembled using Megahit v1.2.x^(53)^ with the kmers 21, 29, 39, 59, 79, 99, 119 and 141 for a minimum contig-length of 100 bp; the assemblies were quality-checked by MetaQUAST^54^; >100-bp-long contigs were searched for ORFs/genes encoding >30-amino-acids-long peptides, using MetaGeneMark^55^.

For the genomes of pure-culture isolates, high quality reads were assembled using SPAdes 3.13.0^(56)^, with kmers 21, 33, 55, 77, 99 and 127, and minimum coverage cut-off 35X. Statistics for >200-bp-long contigs was generated from the assemblies using QUAST^57^ and viewed using Bandage^58^; <200-bp-long contigs were filtered out from the final assemblies. The whole genome sequences were deposited to the GenBank and annotated using Prokaryotic Genome Annotation Pipeline (PGAP located at https://www.ncbi.nlm.nih.gov/genome/annotation_prok/) of National Center for Biotechnology Information (NCBI), USA. Raw reads from the individual metagenomic or metatranscriptomic datasets were mapped onto the individual assembled whole genome sequences, or the manually-curated aerobic-metabolism-related gene-catalogs obtained from the individual genome annotations, using Bowtie2 v.2.3.4.3^(52)^ in default local (sensitive) alignment mode.

The rRNA-sequence-free metatranscriptomic dataset was assembled using the python script rnaspades.py, available within SPAdes 3.13.0^(56)^, with default parameters. ORFs/genes encoding continuous stretches of minimum 30 amino acids were predicted in contigs longer than 100 bp using Prodigal v2.6.3^(59)^.

Gene-catalogs obtained from the individual metagenomes or the metatranscriptome were functionally annotated by searching against EggNOG v5.0 database (http://eggnog5.embl.de/download/eggnog_5.0/) with EggNOG-mapper (http://beta-eggnogdb.embl.de/#/app/emapper) using HMMER algorithm. Putative protein sequence catalogs obtained for the individual genomes via PGAP annotation were re-annotated by searching against EggNOG v5.0 database with EggNOG-mapper using HMMER algorithm. Enzymes involved in aerobic respiration, aerobic methane/sulfur oxidation and perchlorate reduction were screened manually on the basis of their KEGG Orthology numbers^60^.

### MPN of aerobic chemoorganoheterotrophs/chemolithoautotrophs

MPN of viable cells of aerobic, chemoorganoheterotrophs and sulfur-chemolithoautotrophs in the individual sediment-samples were calculated, as described previously^61^, using ten-fold dilution series (and three tubes per dilution) of aerobic slurry-cultures in LB (pH 7.0) and ASWT (pH 7.5) broths respectively (details given in Supplementary Methods). Growth responses observed in an MPN-tube series were tallied with a standard MPN Table^61^ (www.jlindquist.net) to get the MPN of the corresponding metabolic-type g^−1^ sediment-sample.

### Enrichment, isolation and identification of aerobic bacterial strains

To test the presence of viable aerobic microorganisms in ASOMZ sediment, microbial consortia were enriched aerobically from the 275 cmbsf sample of SSK42/6 using the following broth media: LB, R2A (pH 7.0), ASWT, MSTY (pH 7.0) and ASWTY (pH 7.5). ASWT^62^ and ASWTY both contained 10 mM Na_2_S_2_O_3_.5H_2_O while MSTY^63^ contained 20 mM Na_2_S_2_O_3_.5H_2_O; MSTY and ASWTY contained 500 mg yeast extract L^−1^. Detailed compositions of all the five media are given in Supplementary Methods. 5% (w/v) sediment-samples were added to the individual broth media contained in cotton-plugged conical flasks having 60% headspaces filled with air; these flasks were incubated aerobically at 15°C on a rotary shaker (150 rpm). Turbidity of cultures and production of sulfuric acid from thiosulfate^62,63^ were taken as signs of microbial growth in LB/R2A and ASWT/MSTY/ASWTY respectively. After phenol red indicator present in the media turned yellow due to production of sulfuric acid from thiosulfate, aerobic bacterial strains were isolated from the enrichments in ASWT, MSTY and ASWTY, via serial dilution, spread-plating onto corresponding agar media, aerobic incubation at 15°C, and then iterative dilution streaking till all colonies in an isolation plate looked similar^63^. Representative colonies from individual pure-plates were designated as strains and maintained in their respective isolation-media - only the *Methylophaga* strains, despite being isolated in ASWT, were maintained in ASW supplemented with 0.3% (v/v) methanol (ASWM) because their growth vigor in ASWT waned after six consecutive sub-cultures. Strains were classified down to the lowest identifiable taxonomic category, as described previously^63^, on the basis of their 16S rRNA gene sequence similarities with validly-published species (http://www.bacterio.net/).

### Tests for anaerobic/fermentative growth/survival

Isolates were tested for anaerobic growth in their respective maintenance media (Table 1) supplemented with NaNO_3_ (4 mM)^64^, Fe_2_O_3_ (125 mM)^65^, MnO_2_ (1 mM)^66^, Na_2_SO_4_ (10 mM)^67^, (CH_3_)_2_SO (56 mM)^68^ and (CH_3_)_3_NO (27 mM)^68^ as electron-acceptors, provided both as mixture of all six compounds or as single respiratory substrate. Isolates were also tested for anaerobic growth in their respective maintenance media supplemented with humic acids (17 mg mL^−1^)^22^, or humic acids (17 mg mL^−1^) and Fe_2_O_3_ (40 mM) in combination^22^. Isolates which died in these culture conditions were tested for fermentative growth/survival. The *Cereibacter*, ASWTY-isolated-*Halomonas*, *Pseudomonas* and *Sulfitobacter* isolates could not grow aerobically/anaerobically in fermentative medium generally used for lactobacilli^69^; so they were provided pyruvate (5 g L^−1^) as fermentative substrate. Since fructose is the only multicarbon compound utilized by known *Methylophaga* species^24^, the current isolate belonging to this genus was tested for fermentative growth in ASWM medium supplemented with 0.3% (w/v) fructose. Under anaerobic condition all *Halothiobacillus* species lose their obligate sulfur-chemolithotrophic attributes and reportedly perform heterolactic fermentation of stored polyglucose^25^; the present *Halothiobacillus* isolate (SB14A), therefore, was subjected to prolonged incubation in ASWT to test the potential role of such processes in the anoxic survival of this strain.

All components of the above media were mixed, supplemented with 0.5 g L^−1^ sodium thioglycolate (to eliminate O_2_) and 0.1 mg L^−1^ resazurin (to indicate presence of O_2_), and then autoclaved in screw-capped bottles. Methanol was added via filter sterilization after the screw-capped bottles were opened inside the H35 Hypoxystation preset to zero O_2_. Likewise, phosphate, thiosulfate and pyruvate were dissolved in O_2_-free water, filter-sterilized, and added to the relevant media, inside the anaerobic workstation. 100 mL aliquots of individual O_2_-free test-media were dispensed into narrow-mouth and fixed-joint Erlenmeyer flasks, following which cells harvested from 1 mL aerobically-grown mid-log phase seed cultures were added to them; finally the flasks were covered with sleeve stopper septa and incubated at 15°C; all these steps were carried out within the anaerobic workstation. Viable cell count in an anaerobic/fermentative culture at desired time-points of incubation (including 0 h) was determined by counting the colony forming units (CFU) present. To determine CFU-count, 1 mL of the liquid culture was taken out, its various dilution grades plated in triplicates onto agar plates of the corresponding medium (these were done inside the anaerobic workstation), and finally incubated aerobically (outside the workstation). Colony-counts in the different dilution-plates were multiplied by their dilution factors, summed-up, and averaged to get the CFU count mL^−1^ of the test-culture.

### Enrichment of perchlorate-respiring consortia (PRBC) and their co-culture with *Halothiobacillus* SB14A

2 g each of the different sediment-samples were added (inside the H35 Hypoxystation preset at zero O_2_) to separate 40 mL batches of O_2_-free PBBB medium (pH 7.0; composition given in Supplementary Methods)^70^ contained in fixed-joint Erlenmeyer flasks and supplemented with 0.5 g L^−1^ sodium thioglycolate and 0.1 mg L^−1^ resazurin. All the flasks were covered with sleeve stopper septa and incubated anaerobically at 15°C for 45 days, following which 5% inocula were transferred to fresh batches of 40 mL PBBB media. After 7 day incubation of the sub-cultures, the four enrichment sets which exhibited highest O_2_ and/or biomass production were selected for anaerobic co-culture with SB14A. Accordingly, from the 7-day-old PRBC sub-cultures corresponding to 120 and 260 cmbsf of SSK42/5, and 135 and 275 cmbsf of SSK42/6, 10% inocula were added individually, alongside 1% inoculum from a mid-log phase culture of *Halothiobacillus* SB14A, to 40 mL of an O_2_-free, PBBB-ASWT consensus medium (pH 7.0; composition given in Supplementary Methods) that contained energy/electron/carbon sources suitable for perchlorate-respirers as well as *Halothiobacillus*, but perchlorate as sole electron acceptor (O_2_-containing version of this consensus medium was found to support aerobic growth of SB14A, thereby proving its non-toxicity towards this chemolithoautotroph; see Supplementary Table 31). The consensus medium with or without perchlorate was dispensed into 100 mL fixed-joint Erlenmeyer flasks, inoculated with washed cell pellets of both types of cultures or only SB14A harvested according to the inoculums percentages stated above, and covered with sleeve stopper septa inside the anaerobic workstation. The cultures were incubated anaerobically at 15°C for 60 days, and *Halothiobacillus* CFU present mL^−1^ determined, as stated above, at appropriate time-intervals (including 0 h) by dilution plating in ASWT-agar followed by aerobic incubation at 15°C. Three discrete colonies from each CFU-counting plate were checked by 16S rRNA gene sequence analysis to corroborate their *Halothiobacillus* identity.

### Analytical methods

To corroborate sulfur-chemolithotrophic/mixotrophic growth of pure-cultures/microbial consortia, concentrations of dissolved thiosulfate and sulfate in the spent media were measured by iodometric titration, and gravimetric precipitation with BaCl_2_, respectively^64^. Potential concentration of dissolved perchlorate in the sediment pore-waters were determined by ion chromatography using an Eco IC (Metrohm AG, Switzerland) equipped with a conductivity detector (Metrohm, IC detector 1.850.9010). Chemical suppression was used for this purpose, while separation was carried using a Metrosep A Supp5 - 250/4.0 (6.1006.530) anion exchange column (Metrohm AG). A mixture of 1.0 mM sodium hydrogen carbonate and 3.2 mM sodium carbonate was used as the eluent; 100 mM sulfuric acid was used as the regenerant; injection volume was 100 µL, and flow rate 0.7 mL min^−1^. Samples were diluted 1000-fold with de-ionized water (Siemens, <0.06 μS), and filtered using 0.22 µm hydrophilic polyvinylidene fluoride membranes (Merck Life Science Private Limited, India), prior to analysis. Sigma-Aldrich (USA) standard chemicals were used to prepare the calibration curve for quantification; overall sample reproducibility was ±0.2 ppm.

### Code availability

All data analysis codes used in this study are in the published domain, and have been appropriately cited in the text.

### Data availability

All nucleotide sequence data have been deposited in NCBI Sequence Read Archive (SRA) or GenBank under the BioProject accession number PRJNA309469: (i) the whole metagenome shotgun sequence datasets have the Run accession numbers SRR3646127 through SRR3646132, SRR3646144, SRR3646145, SRR3646147, SRR3646148, SRR3646150 through SRR3646153, SRR3646155 through SRR3646158, SRR3646160 through SRR3646165; SRR3570036, SRR3570038, SRR3577067, SRR3577068, SRR3577070, SRR3577071, SRR3577073, SRR3577076, SRR3577078, SRR3577079, SRR3577081, SRR3577082, SRR3577086, SRR3577087, SRR3577090, SRR3577311, SRR3577337, SRR3577338, SRR3577341, SRR3577343, SRR3577344, SRR3577345, SRR3577347, SRR3577349, SRR3577350, SRR3577351; SRR3872933, SRR3872934, SRR3884351, SRR3884355, SRR3884357, SRR3884359, SRR3884468, SRR3884472, SRR3884479, SRR3884488, SRR3884538, SRR3884540, SRR3884542, SRR3884544, SRR3884546 through SRR3884548, SRR3884552, SRR3884553 and SRR3884554; (ii) the whole metatranscriptome sequence dataset has the Run accession number SRR7991972; (iii) the whole genome sequences have the GenBank accession numbers SWAW01000000, SSXS01000000, SZNL01000000, SSXT01000000, SWAV01000000 and SWAU01000000.

## Acknowledgements

Financial support for conducting the microbiological studies was provided by given by Bose Institute via internal faculty grants and Earth System Science Organization, Ministry of Earth Sciences (MoES), Government of India (GoI) via grant number MoES/36/00IS/Extra/19/2013. We thank the Director CSIR-National Institute of Oceanography for facilitating the geochemical studies and the research cruise SSK42 for acquisition of sediment cores. MoES (GAP2303) also funded the research cruise. All the support received from the CSIR-NIO Ship Cell members and the crew members of SSK42 is gratefully acknowledged. S.B. and M.A. received fellowship from Bose Institute. C.R. and MJR got fellowship from University Grants Commission, GoI. SM got fellowship from Department of Science and Technology, GoI. J.S., S.F., R.R. and P.P received fellowships from Council of Scientific and Industrial Research, GoI. N.M. got fellowship from Science and Engineering Research Board, GoI, under the grant EMR/2016/002703. We thank Werner Liesack (Max Planck Institute for Terrestrial Microbiology, Germany), Fumio Inagaki (Japan Agency for Marine-Earth Science and Technology, Japan) and Subhra Kanti Mukhopadhyay (University of Burdwan, India) for useful discussions.

## Author contributions

W.G. conceived the study, designed the experiments, interpreted the results and wrote the paper. S.B. anchored the whole microbiological study, performed the experiments, and analyzed and curated the data. A.M. led the mission SSK42 and all geochemical investigations therein. R.C. and A.M. participated in result interpretation and made intellectual contributions. C.R., M.J.R., J.S., T.M. and M.A. performed bioinformatic and microbiological data analyses. S.M., R.R., N.M., P.P. and P.K.H. performed microbiological experiments. S.F. and A.P. performed geochemical experiments. All authors read and vetted the manuscript.

## Additional information

Supplementary Information and Dataset accompany the online version of the paper.

## Competing interest

The authors declare no competing interest.

## References

1. Wright, J. J., Konwar, K. M. & Hallam, S. J. Microbial ecology of expanding oxygen minimum zones. Nat. Rev. Microbiol. 10, 381 (2012).

2. Fernandes, S. et al. Enhanced carbon-sulfur cycling in the sediments of Arabian Sea oxygen minimum zone center. Sci. Rep. 8, 8665 (2018).

3. Maltby, J., Sommer, S., Dale, A. W. & Treude, T. Microbial methanogenesis in the sulfate-reducing zone of surface sediments traversing the Peruvian margin. Biogeosciences 13, 283–299 (2016).

4. Ulloa, O., Canfield, D. E., DeLong, E. F., Letelier, R. M., & Stewart, F. J. Microbial oceanography of anoxic oxygen minimum zones. Proc. Natl. Acad. Sci. USA 109, 15996–16003 (2012).

5. Bertagnolli, A. D. & Stewart, F. J. Microbial niches in marine oxygen minimum zones. Nat. Rev. Microbiol. 16, 723–729 (2018).

6. Lam, P. et al. Revising the nitrogen cycle in the Peruvian oxygen minimum zone. Proc. Natl. Acad. Sci. USA 106, 4752–4757 (2009).

7. Revsbech, N. P. et al. Determination of ultra-low oxygen concentrations in oxygen minimum zones by the STOX sensor. Limnol. Oceanogr-Meth. 7, 371–381 (2009).

8. Canfield, D. E. et al. A cryptic sulfur cycle in oxygen-minimum–zone waters off the Chilean coast. Science 330, 1375–1378 (2010).

9. Stolper, D. A., Revsbech, N. P. & Canfield, D. E. Aerobic growth at nanomolar oxygen concentrations. Proc. Natl. Acad. Sci. USA 107, 18755–18760 (2010).

10. Füssel, J. et al. Nitrite oxidation in the Namibian oxygen minimum zone. ISME J. 6, 1200–1209 (2012).

11. Riedel, T. E., Berelson, W. M., Nealson, K. H. & Finkel, S. E. Oxygen consumption rates of bacteria under nutrient-limited conditions. Appl. Environ. Microb. 79, 4921–4931 (2013).

12. Kalvelage, T. et al. Aerobic microbial respiration in oceanic oxygen minimum zones. PloS one 10, e0133526 (2015).

13. Garcia-Robledo, E. et al. Cryptic oxygen cycling in anoxic marine zones. Proc. Natl. Acad. Sci. USA 114, 8319–8324 (2017).

14. Breuer, E. R. et al. Sedimentary oxygen consumption and microdistribution at sites across the Arabian Sea oxygen minimum zone (Pakistan margin). Deep-Sea. Res. Pt II. 56, 296–304 (2009).

15. Cavan, E. L., Trimmer, M., Shelley, F. & Sanders, R. Remineralization of particulate organic carbon in an ocean oxygen minimum zone. Nat. Commun. 8, 14847 (2017).

16. Jessen, G. L. et al. Hypoxia causes preservation of labile organic matter and changes seafloor microbial community composition (Black Sea). Sci. Adv. 3, e1601897 (2017).

17. Acharya, S. S. & Panigrahi, M. K. Eastward shift and maintenance of Arabian Sea oxygen minimum zone: Understanding the paradox. Deep Sea Res. Part I Oceanogr. Res. Pap. 115, 240–252 (2016).

18. Canfield, D. E., Kristensen, E. & Thamdrup, B. in Aquatic geomicrobiology 656 (Academic Press, 2005).

19. López, N. I. & Duarte, C. M. Dimethyl sulfoxide (DMSO) reduction potential in Mediterranean seagrass (Posidonia oceanica) sediments. J. Sea Res. 51, 11–20 (2004).

20. Jørgensen, B. B. & Kasten, S. in Marine Geochemistry (ed. Schulz, H. D. & Zabel, M.) 271–309 (Springer, Berlin, 2006).

21. Lidbury, I., Murrell, J. C. & Chen, Y. Trimethylamine N-oxide metabolism by abundant marine heterotrophic bacteria. Proc. Natl. Acad. Sci. USA 111, 2710–2715 (2014).

22. Benz, M., Schink, B. & Brune, A. Humic acid reduction by Propionibacterium freudenreichii and other fermenting bacteria. Appl. Environ. Microbiol. 64, 4507–4512 (1998).

23. He, S., Lau, M. P., Linz, A. M., Roden, E. E. & McMahon, K. D. Extracellular electron transfer may be an overlooked contribution to pelagic respiration in humic-rich freshwater lakes. mSphere 4, e00436–18 (2019).

24. Janvier, M., Frehel, C., Grimont, F. & Gasser, F. Methylophaga marina gen. nov., sp. nov. and Methylophaga thalassica sp. nov., marine methylotrophs. Int. J. Syst. Evol. Microbiol. 35, 131–139 (1985).

25. Beudeker, R., Boer, W. & Kuenen, J. Heterolactic fermentation of intracellular polyglucose by the obligate chemolithotroph *Thiobacillus neapolitanus* under anaerobic conditions. FEMS Microbiol. Lett. 12, 337–342 (1981).

26. Blazewicz, S. J., Barnard, R. L., Daly, R. A. & Firestone, M. K. Evaluating rRNA as an indicator of microbial activity in environmental communities: limitations and uses. The ISME J. 7, 2061–2068 (2013).

27. Johnson, S. S. et al. Ancient bacteria show evidence of DNA repair. Proc. Natl. Acad. Sci. USA 104, 14401–14405 (2007).

28. Lomstein, B. A., Langerhuus, A. T., D’Hondt, S., Jørgensen, B. B. & Spivack, A. J. Endospore abundance, microbial growth and necromass turnover in deep sub-seafloor sediment. Nature 484, 101–104 (2012).

29. Kempes, C. P., van Bodegom, P. M., Wolpert, D., Libby, E., Amend, J. & Hoehler, T. Drivers of bacterial maintenance and minimal energy requirements. Front. Microbiol. 8, 1–10 (2017).

30. Jiang, Y., Xiong, X., Danska, J. & Parkinson, J. Metatranscriptomic analysis of diverse microbial communities reveals core metabolic pathways and microbiome-specific functionality. Microbiome 4, 2 (2016).

31. Bardiya, N. & Bae, J-H. Dissimilatory perchlorate reduction: a review. Microbiol. Res. 166, 237–254. (2011).

32. Parker, G. A. & Smith, J. M. Optimality theory in evolutionary biology. Nature, 348, 27–33 (1990).

33. Moran, et al. Sizing up metatranscriptomics. The ISM E J. 7, 237–243 (2013).

34. Rauhut, R. & Klug, G. mRNA degradation in bacteria. FEMS Microbiol. Rev. 23, 353–370 (1999).

35. Steglich, C., Lindell, D., Futschik, M., Rector, T., Steen, R. & Chisholm, S. W. Short RNA half-lives in the slow-growing marine cyanobacterium *Prochlorococcus*. Genome Biol. 11, 2–14 (2010).

36. Hundt, S., Zaigler, A., Lange, C., Soppa, J. & Klug, G. Global analysis of mRNA decay in *Halobacterium salinarum* NRC-1 at single-gene resolution using DNA microarrays. J. Bacteriol. 189, 6936–6944 (2007).

37. Steiner, P. A. et al. Highly variable mRNA half‐life time within marine bacterial taxa and functional genes. Environ. Microbiol. (2019).

38. Orsi, W., Biddle, J. F. & Edgcomb, V. Deep sequencing of subseafloor eukaryotic rRNA reveals active fungi across marine subsurface provinces. PLoS One 8, p.e56335 (2013).

39. Ignatov, D. V., Salina, E. G., Fursov, M. V., Skvortsov, T. A., Azhikina, T. L. & Kaprelyants, S. Dormant non-culturable *Mycobacterium tuberculosis* retains stable low-abundant mRNA. BMC genomics 16, 954 (2015).

40. Orsi, W. D., Edgcomb, V. P., Christman, G. D. & Biddle, J. F. Gene expression in the deep biosphere. Nature 499, 205–208 (2013).

41. Thureborn, P., Franzetti, A., Lundin, D. & Sjöling, S. Reconstructing ecosystem functions of the active microbial community of the Baltic Sea oxygen depleted sediments. PeerJ 4, e1593 (2016).

42. Coates, J. D. & Achenbach, L. A. Microbial perchlorate reduction: rocket-fueled metabolism. Nat. Rev. Microbiol. 2, 569–580 (2004).

43. Carlström, C. I. et al. Physiological and genetic description of dissimilatory perchlorate reduction by the novel marine bacterium *Arcobacter sp*. strain CAB. MBio 4, e00217–13 (2013).

44. Carlström, C. I. et al. Phenotypic and genotypic description of *Sedimenticola selenatireducens* strain CUZ, a marine (per) chlorate-respiring gammaproteobacterium, and its close relative the chlorate-respiring *Sedimenticola* strain NSS. Appl. Environ. Microbiol. 81, 2717–2726 (2015).

45. Stepanov, V. G., Xiao, Y., Lopez, A. J., Roberts, D. J. & Fox, G. E. Draft genome sequence of *Marinobacter sp*. strain P4B1, an electrogenic perchlorate-reducing strain isolated from a long-term mixed enrichment culture of marine bacteria. Genome Announc. 4, e01617–15 (2016).

46. Kamp, A., Høgslund, S., Risgaard-Petersen, N. & Stief, P. Nitrate storage and dissimilatory nitrate reduction by eukaryotic microbes. Front. Microbiol. 6, 1492 (2015).

47. Olsson, I. U. in. Radiocarbon Variations and Absolute Chronology, (eds Olsson, I. U.) 17 (Nobel Symposium, 12th Proc., John Wiley & Sons, New York, 1970).

48. Fairbanks, R. G. Radiocarbon calibration curve spanning 0 to 50,000 years BP based on paired ^230^Th/^234^U/^238^U and ^14^C dates on pristine corals. Quat. Sci. Rev. 24, 1781–1796 (2005).

49. Stuiver, M. & Polach, H. A. Discussion Reporting of ^14^C data. Radiocarbon 19, 355–363 (1977).

50. Ghosh, W. et al. Resilience and receptivity worked in tandem to sustain a geothermal mat community amidst erratic environmental conditions. Sci. Rep. 5, 12179 (2015).

51. Quast, C. et al. The SILVA ribosomal RNA gene database project: improved data processing and web-based tools. Nucleic Acids Res. 41, D590–D596 (2013).

52. Langmead, B. & Salzberg, S. L. Fast gapped-read alignment with Bowtie2. Nat. Methods 9, 357–359 (2012).

53. Li, D., Liu, C. M., Luo, R., Sadakane, K. & Lam, T. W. MEGAHIT: an ultra-fast single-node solution for large and complex metagenomics assembly via succinct de Bruijn graph. Bioinformatics 31, 1674–1676 (2015).

54. Mikheenko, A., Saveliev, V. & Gurevich, A. MetaQUAST: evaluation of metagenome assemblies. Bioinformatics 32, 1088–1090 (2016).

55. Zhu, W., Lomsadze, A., Borodovsky, M. Ab initio gene identification in metagenomic sequences. Nucleic Acids Res. 38, e132 (2010).

56. Nurk, S., et al. Assembling single-cell genomes and mini-metagenomes from chimeric MDA products. J. Comput. Biol. 20, 714–737 (2013).

57. Gurevich, A., Saveliev, V., Vyahhi, N. & Tesler, G. QUAST: quality assessment tool for genome assemblies. Bioinformatics 29, 1072–1075 (2013).

58. Wick, R. R., Schultz, M. B., Zobel, J. & Holt, K. E. Bandage: interactive visualization of de novo genome assemblies. Bioinformatics 31, 3350–3352 (2015).

59. Hyatt, D., Chen, G. L., Locascio, P.F., Land, M. L., Larimer, F. W. & Hauser, L. J. Prodigal: prokaryotic gene recognition and translation initiation site identification. BMC Bioinformatics 11, 119 (2010).

60. Kanehisa, M., Sato, Y., Kawashima, M., Furumichi, M. & Tanabe, M. KEGG as a reference resource for gene and protein annotation. Nucleic Acids Res. 44, D457–462 (2016).

61. Sutton, S. The most probable number method and its uses in enumeration, qualification, and validation. Microbiol. Top. J. Validation Tech. 16, 35–38 (2010).

62. Alam, M., Pyne, P., Mazumdar, A., Peketi, A. & Ghosh, W. Kinetic enrichment of ^34^S during proteobacterial thiosulfate oxidation and the conserved role of SoxB in S-S bond breaking. Appl. Environ. Microbiol. 79, 4455–4464 (2013).

63. Ghosh, W. & Roy, P. Mesorhizobium thiogangeticum sp. nov., a novel sulfur-oxidizing chemolithoautotroph from rhizosphere soil of an Indian tropical leguminous plant. Int. J. Syst. Evol. Microbiol. 56, 91–97 (2006).

64. Straub, K. L. & Buchholz-Cleven, B. E. Enumeration and detection of anaerobic ferrous iron-oxidizing, nitrate-reducing bacteria from diverse European sediments. Appl. Environ. Microbiol. 64, 4846–4856 (1998).

65. Lovley, D. R. & Phillips, E. J. Organic matter mineralization with reduction of ferric iron in anaerobic sediments. Appl. Environ. Microbiol. 51, 683–689 (1986).

66. Myers, C. R. & Nealson, K. H. Bacterial manganese reduction and growth with manganese oxide as the sole electron acceptor. Science 240, 1319–1321 (1988).

67. So, C. M. & Young, L. Isolation and characterization of a sulfate-reducing bacterium that anaerobically degrades alkanes. Appl. Environ. Microbiol. 65, 2969–2976 (1999).

68. Oren, A. & Trüper, H. G. Anaerobic growth of halophilic archaeobacteria by reduction of dimethylsulfoxide and trimethylamine N-oxide. FEMS Microbiol. Lett. 70, 33–36 (1990).

69. Mercier, P., Yerushalmi, L., Rouleau, D. & Dochain, D. Kinetics of lactic acid fermentation on glucose and corn by Lactobacillus amylophilus. J. Chem. Technol. Biotechnol. 55, 111–121 (1992).

70. Balk, M., van Gelder, T., Weelink, S. A. & Stams, A. J. (Per)chlorate reduction by the thermophilic bacterium *Moorella perchloratireducens* sp. nov., isolated from underground gas storage. Appl. Environ. Microbiol. 74, 403–409 (2008).

