## Supplementary Information for "Metabolically-active obligate aerobes in anoxic (sulfidic) marine sediments"

Running Title: Strictly-aerobic life in anoxic sediments

**Contents**

**Supplementary Figure**

**Supplementary Figure 1.** Schematic diagram showing the locations of the currently-explored sediment-cores SSK42/5, SSK42/6 and SSK42/9, in the context of the oxygen minimum zone and the other SSK42 cores studied from this region previously.

**Supplementary Tables**

**Supplementary Tables 1 and 2.** Sediment-depths of SSK42/5 and 6 explored for metagenome analysis, respectively. Sequence accession numbers for the individual sediment-samples are also given.

**Supplementary Table 3.** Number of aerobic-respiration-related genes identified within the metagenome assemblies obtained for the individual sediment-samples of SSK42/5 and 6. Since this Supplementary Table is more than one page long it has been provided as an Excel sheet named Supplementary Table 3, within the Excel Workbook named Supplementary Dataset.

**Supplementary Table 4.** Number of aerobic methane-oxidation-related genes identified within the metagenome assemblies obtained for the individual sediment-samples of SSK42/5 and 6. Since this Supplementary Table is more than one page long it has been provided as an Excel sheet named Supplementary Table 4, within the Excel Workbook named Supplementary Dataset.

**Supplementary Table 5.** Number of aerobic sulfur-oxidation-related genes identified within the metagenome assemblies obtained for the individual sediment-samples of SSK42/5 and 6. Since this Supplementary Table is more than one page long it has been provided as an Excel sheet named Supplementary Table 5, within the Excel Workbook named Supplementary Dataset.

**Supplementary Table 6.** Sediment-depths of SSK42/9 explored for metagenome analysis. Pore-water sulfide concentrations, and metagenome sequence accession numbers, are also given for the individual sediment-samples explored.

**Supplementary Table 7.** Number of aerobic-respiration-related genes identified within the metagenome assemblies obtained for the individual sediment-samples of SSK42/9. Since this Supplementary Table is more than one page long it has been provided as an Excel sheet named Supplementary Table 7, within the Excel Workbook named Supplementary Dataset.

**Supplementary Table 8.** Number of aerobic methane-oxidation-related genes identified within the metagenome assemblies obtained for the individual sediment-samples of SSK42/9. Since this Supplementary Table is more than one page long it has been provided as an Excel sheet named Supplementary Table 8, within the Excel Workbook named Supplementary Dataset.

**Supplementary Table 9.** Number of aerobic sulfur-oxidation-related genes identified within the metagenome assemblies obtained for the individual sediment-samples of SSK42/9. Since this Supplementary Table is more than one page long it has been provided as an Excel sheet named Supplementary Table 9, within the Excel Workbook named Supplementary Dataset.

**Supplementary Tables 10 and 11**. Most probable number of aerobic, chemoorganohetarotrophic microorganisms along the explored sediment-depths of SSK42/5 and 6, respectively.

**Supplementary Tables 12 and 13**. Most probable number of aerobic, chemolithoautotrophic microorganisms along the explored sediment-depths of SSK42/5 and 6 respectively.

**Supplementary Table 14.** Aerobic growth of the nine representative isolates from 275 cmbsf of SSK42/6 in media lacking/containing the alternative electron acceptors that were subsequently used in anaerobic growth experiments. Since this Supplementary Table is more than one page long it has been provided as an Excel sheet named Supplementary Table 14, within the Excel Workbook named Supplementary Dataset.

**Supplementary Table 15.** Results of anaerobic incubation of the nine representative isolates from 275 cmbsf of SSK42/6 in media containing different alternative electron acceptors. Since this Supplementary Table is more than one page long it has been provided as an Excel sheet named Supplementary Table 15, within the Excel Workbook named Supplementary Dataset.

**Supplementary Table 16.** General features of the assembled genomes of the 6 obligately aerobic strains isolated from 275 cmbsf of SSK42/6, together with the percentages of metagenomic reads from individual sediment-samples of SSK42/5, 6 and 9 that matched with sequences from each genome. Since this Supplementary Table is more than one page long it has been provided as an Excel sheet named Supplementary Table 16, within the Excel Workbook named Supplementary Dataset.

**Supplementary Table 17.** Aerobic-metabolism-related genes of *Cereibacter changlaensis* MTCC 12557, and the numbers of metatranscriptomic read-pairs from 275 cmbsf of SSK42/6 that matched concordantly with the individual genes of this catalog. Since this Supplementary Table is more than one page long it has been provided as an Excel sheet named Supplementary Table 17, within the Excel Workbook named Supplementary Dataset.

**Supplementary Table 18.** Aerobic-metabolism-related genes of *Halomonas* sp. MCC 3301, and the numbers of metatranscriptomic read-pairs from 275 cmbsf of SSK42/6 that matched concordantly with the individual genes of this catalog. Since this Supplementary Table is more than one page long it has been provided as an Excel sheet named Supplementary Table 18, within the Excel Workbook named Supplementary Dataset.

**Supplementary Table 19.** Aerobic-metabolism-related genes of *Halothiobacillus* sp. SB14A, and the numbers of metatranscriptomic read-pairs from 275 cmbsf of SSK42/6 that matched concordantly with the individual genes of this catalog. Since this Supplementary Table is more than one page long it has been provided as an Excel sheet named Supplementary Table 19, within the Excel Workbook named Supplementary Dataset.

**Supplementary Table 20.** Aerobic-metabolism-related genes of *Methylophaga* sp. MTCC 12599, and the numbers of metatranscriptomic read-pairs from 275 cmbsf of SSK42/6 that matched concordantly with the individual genes of this catalog. Since this Supplementary Table is more than one page long it has been provided as an Excel sheet named Supplementary Table 20, within the Excel Workbook named Supplementary Dataset.

**Supplementary Table 21.** Aerobic-metabolism-related genes of *Pseudomonas bauzanensis* MTCC 12600, and the numbers of metatranscriptomic read-pairs from 275 cmbsf of SSK42/6 that matched concordantly with the individual genes of this catalog. Since this Supplementary Table is more than one page long it has been provided as an Excel sheet named Supplementary Table 21, within the Excel Workbook named Supplementary Dataset.

**Supplementary Table 22.** Aerobic-metabolism-related genes of *Sulfitobacter* sp. MCC 3606, and the numbers of metatranscriptomic read-pairs from 275 cmbsf of SSK42/6 that matched concordantly with the individual genes of this catalog. Since this Supplementary Table is more than one page long it has been provided as an Excel sheet named Supplementary Table 22, within the Excel Workbook named Supplementary Dataset.

**Supplementary Table 23.** Core-metabolism-related genes of *Halothiobacillus* sp. SB14A, and the numbers of metatranscriptomic read-pairs from 275 cmbsf of SSK42/6 that matched concordantly with the individual genes of this catalog. Since this Supplementary Table is more than one page long it has been provided as an Excel sheet named Supplementary Table 23, within the Excel Workbook named Supplementary Dataset.

**Supplementary Table 24.** Core-metabolism-related genes of *Cereibacter changlaensis* MTCC 12557, and the numbers of metatranscriptomic read-pairs from 275 cmbsf of SSK42/6 that matched concordantly with the individual genes of this catalog. Since this Supplementary Table is more than one page long it has been provided as an Excel sheet named Supplementary Table 24, within the Excel Workbook named Supplementary Dataset.

**Supplementary Table 25.** Core-metabolism-related genes of *Halomonas* sp. MCC 3301, and the numbers of metatranscriptomic read-pairs from 275 cmbsf of SSK42/6 that matched concordantly with the individual genes of this catalog. Since this Supplementary Table is more than one page long it has been provided as an Excel sheet named Supplementary Table 25, within the Excel Workbook named Supplementary Dataset.

**Supplementary Table 26.** Core-metabolism-related genes of *Pseudomonas bauzanensis* MTCC 12600, and the numbers of metatranscriptomic read-pairs from 275 cmbsf of SSK42/6 that matched concordantly with the individual genes of this catalog. Since this Supplementary Table is more than one page long it has been provided as an Excel sheet named Supplementary Table 26, within the Excel Workbook named Supplementary Dataset.

**Supplementary Table 27.** Core-metabolism-related genes of *Sulfitobacter* sp. MCC 3606, and the numbers of metatranscriptomic read-pairs from 275 cmbsf of SSK42/6 that matched concordantly with the individual genes of this catalog. Since this Supplementary Table is more than one page long it has been provided as an Excel sheet named Supplementary Table 27, within the Excel Workbook named Supplementary Dataset.

**Supplementary Table 28.** Core-metabolism-related genes of *Methylophaga* sp. MTCC 12599, and the numbers of metatranscriptomic read-pairs from 275 cmbsf of SSK42/6 that matched concordantly with the individual genes of this catalog. Since this Supplementary Table is more than one page long it has been provided as an Excel sheet named Supplementary Table 28, within the Excel Workbook named Supplementary Dataset.

**Supplementary Table 29.** Number of perchlorate-respiration-related genes identified within the metagenome assemblies obtained for the individual sediment-samples of SSK42/5 and 6. Since this Supplementary Table is more than one page long it has been provided as an Excel sheet named Supplementary Table 29, within the Excel Workbook named Supplementary Dataset.

**Supplementary Table 30.** Number of perchlorate-respiration-related genes identified within the metagenome assemblies obtained for the individual sediment-samples of SSK42/9. Since this Supplementary Table is more than one page long it has been provided as an Excel sheet named Supplementary Table 30, within the Excel Workbook named Supplementary Dataset.

**Supplementary Table 31.** Growth behavior of *Halothiobacillus* sp. SB14A in co-cultures with or without perchlorate-respiring bacterial consortia, in a consensus medium supplemented with or without perchlorate and containing energy, electron and carbon sources suitable for both *Halothiobacillus* and perchlorate-respirers.

**Supplementary Table 32.** (Per)chlorate reductase genes identified within the contigs assembled from the metatranscriptomic sequence data obtained from the 275 cmbsf sediment-sample of SSK42/6.

**Supplementary Methods**

Details of the methods used in determining Most Probable Number (MPN) of aerobic bacteria, and metagenome/genome/metatranscriptome sequencing.

**Supplementary References**

Papers cited in Supplementary Methods are given.

**Supplementary Figure**

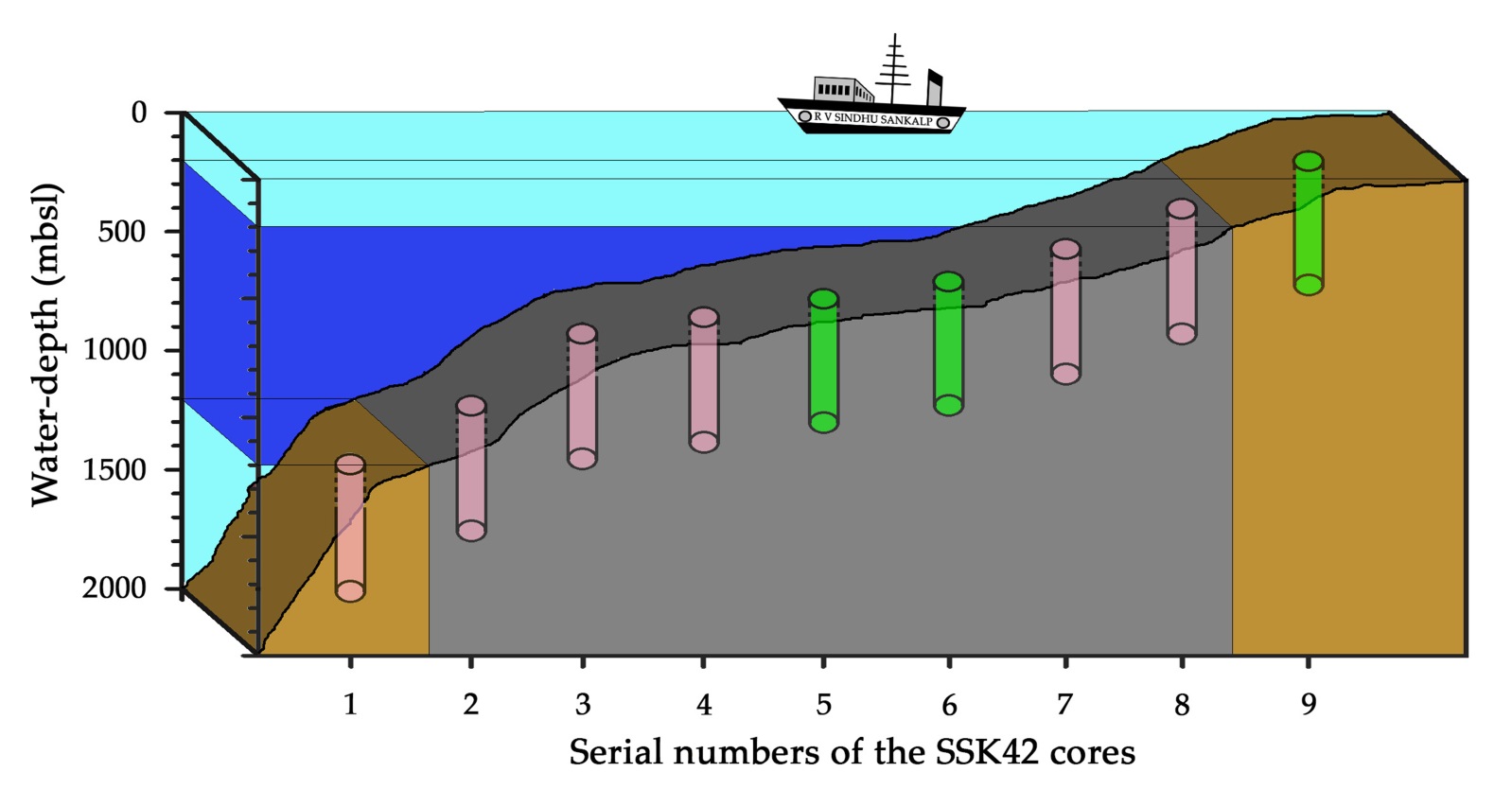

**Supplementary Figure 1.** **Schematic diagram showing the locations of the currently-explored sediment-cores SSK42/5, SSK42/6 and SSK42/9 in the context of the oxygen minimum zone and the other SSK42 cores studied from this region previously** **Fernandes et al. [*Sci*. *Rep*. 8, 8665 (2018); see reference number 2 of the main article file].** Water-depth is plotted in scale along the vertical axis of the diagram, while distances between the cores represented along the horizontal axis are not in scale. The oxygen minimum zone is indicated by blue shade; both the upper and lower oxyclines are indicated by light turquoise shade; sediments horizons underlying the oxygen minimum zone is indicated by gray shade and sediments horizons underlying upper and lower oxyclines are indicated by brown shade.

**Supplementary Tables**

**Supplementary Table 1.** Sediment-depths of SSK42/5 explored for duplicate metagenome sequencing-based structure/function of the community.

| **Sediment-depth explored**  **(in cmbsf)** | **BioSample accession number** | **Sample-fractions** | **Run accession number of the metagenomic sequence dataset** |
| --- | --- | --- | --- |
| 0 | SAMN04442175 | 1st fraction | SRR3646127 |
|  |  | 2nd fraction | SRR3646128 |
| 15 | SAMN04442176 | 1st fraction | SRR3646129 |
|  |  | 2nd fraction | SRR3646130 |
| 45 | SAMN04442177 | 1st fraction | SRR3646131 |
|  |  | 2nd fraction | SRR3646132 |
| 60 | SAMN04442178 | 1st fraction | SRR3646144 |
|  |  | 2nd fraction | SRR3646145 |
| 90 | SAMN04442179 | 1st fraction | SRR3646147 |
|  |  | 2nd fraction | SRR3646148 |
| 120 | SAMN04442180 | 1st fraction | SRR3646150 |
|  |  | 2nd fraction | SRR3646151 |
| 140 | SAMN04442181 | 1st fraction | SRR3646152 |
|  |  | 2nd fraction | SRR3646153 |
| 160 | SAMN04442182 | 1st fraction | SRR3646155 |
|  |  | 2nd fraction | SRR3646156 |
| 190 | SAMN04442183 | 1st fraction | SRR3646157 |
|  |  | 2nd fraction | SRR3646158 |
| 220 | SAMN04442184 | 1st fraction | SRR3646160 |
|  |  | 2nd fraction | SRR3646161 |
| 260 | SAMN04442185 | 1st fraction | SRR3646162 |
|  |  | 2nd fraction | SRR3646163 |
| 295 | SAMN04442186 | 1st fraction | SRR3646164 |
|  |  | 2nd fraction | SRR3646165 |

**Supplementary Table 2.** Sediment-depths of SSK42/6 explored for duplicate metagenome sequencing-based structure/function of the community.

| **Sediment-depth explored**  **(in cmbsf)** | **BioSample accession number** | **Sample-fractions** | **Run accession number of the metagenomic sequence dataset** |
| --- | --- | --- | --- |
| 2 | SAMN04442187 | 1st fraction | SRR3570036 |
|  |  | 2nd fraction | SRR3570038 |
| 30 | SAMN04442189 | 1st fraction | SRR3577067 |
|  |  | 2nd fraction | SRR3577068 |
| 45 | SAMN04442190 | 1st fraction | SRR3577070 |
|  |  | 2nd fraction | SRR3577071 |
| 60 | SAMN04442191 | 1st fraction | SRR3577073 |
|  |  | 2nd fraction | SRR3577076 |
| 75 | SAMN04442192 | 1st fraction | SRR3577078 |
|  |  | 2nd fraction | SRR3577079 |
| 90 | SAMN04442193 | 1st fraction | SRR3577081 |
|  |  | 2nd fraction | SRR3577082 |
| 120 | SAMN04442195 | 1st fraction | SRR3577086 |
|  |  | 2nd fraction | SRR3577087 |
| 135 | SAMN04442196 | 1st fraction | SRR3577090 |
|  |  | 2nd fraction | SRR3577311 |
| 175 | SAMN04442199 | 1st fraction | SRR3577337 |
|  |  | 2nd fraction | SRR3577338 |
| 220 | SAMN04442202 | 1st fraction | SRR3577341 |
|  |  | 2nd fraction | SRR3577343 |
| 250 | SAMN04442204 | 1st fraction | SRR3577344 |
|  |  | 2nd fraction | SRR3577345 |
| 265 | SAMN04442205 | 1st fraction | SRR3577347 |
|  |  | 2nd fraction | SRR3577349 |
| 275 | SAMN04442207 | 1st fraction | SRR3577350 |
|  |  | 2nd fraction | SRR3577351 |

**Supplementary Table 6.** Sediment-depths of SSK42/9 explored for duplicate metagenome sequencing-based structure/function of the community; pore-water sulfide concentrations are also given.

| **Sediment-depth explored**  **(in cmbsf)** | **HS^-^**  **(in µM)** | **BioSample accession number** | **Sample-fractions** | **Run accession number of the metagenomic sequence dataset** |
| --- | --- | --- | --- | --- |
| 0 | 0 | SAMN04442233 | 1st fraction | SRR3872933 |
|  |  |  | 2nd fraction | SRR3872934 |
| 19 | 100 | SAMN04442236 | 1st fraction | SRR3884351 |
|  |  |  | 2nd fraction | SRR3884355 |
| 50 | 700 | SAMN04442238 | 1st fraction | SRR3884357 |
|  |  |  | 2nd fraction | SRR3884359 |
| 115 | 900 | SAMN04442241 | 1st fraction | SRR3884468 |
|  |  |  | 2nd fraction | SRR3884472 |
| 125 | 1100 | SAMN04442242 | 1st fraction | SRR3884479 |
|  |  |  | 2nd fraction | SRR3884488 |
| 145 | 785 | SAMN04442244 | 1st fraction | SRR3884538 |
|  |  |  | 2nd fraction | SRR3884540 |
| 155 | 850 | SAMN04442245 | 1st fraction | SRR3884542 |
|  |  |  | 2nd fraction | SRR3884544 |
| 180 | 1190 | SAMN04442246 | 1st fraction | SRR3884546 |
|  |  |  | 2nd fraction | SRR3884547 |
| 225 | 460 | SAMN04442248 | 1st fraction | SRR3884548 |
|  |  |  | 2nd fraction | SRR3884552 |
| 255 | 960 | SAMN04442250 | 1st fraction | SRR3884553 |
|  |  |  | 2nd fraction | SRR3884554 |

**Supplementary Table 10.** Most probable number of aerobic chemoorganohetarotrophic microorganisms along the explored sediment-depths of SSK42/5.

| **Sediment-depth explored**  **(in cmbsf)** | **Sample dilution grade tested and the number of tubes showing positive growth response therein** | | | | | | | | | | **MPN**  **g^-1^ sediment** |
| --- | --- | --- | --- | --- | --- | --- | --- | --- | --- | --- | --- |
|  | **10^-1^** | **10^-2^** | **10^-3^** | **10^-4^** | **10^-5^** | **10^-6^** | **10^-7^** | **10^-8^** | **10^-9^** | **10^-10^** |  |
| 0 | 3 | 3 | 3 | 2 | 0 | 0 | 0 | 0 | 0 | 0 | 11,000 |
| 15 | 3 | 3 | 3 | 0 | 1 | 0 | 0 | 0 | 0 | 0 | 3,800 |
| 45 | 3 | 3 | 2 | 3 | 0 | 0 | 0 | 0 | 0 | 0 | 2,900 |
| 60 | 3 | 3 | 3 | 0 | 0 | 0 | 0 | 0 | 0 | 0 | 2,400 |
| 90 | 3 | 3 | 1 | 2 | 0 | 0 | 0 | 0 | 0 | 0 | 1,200 |
| 120 | 3 | 3 | 2 | 3 | 1 | 0 | 0 | 0 | 0 | 0 | 3,600 |
| 140 | 3 | 3 | 1 | 2 | 1 | 0 | 0 | 0 | 0 | 0 | 1,500 |
| 160 | 3 | 3 | 2 | 0 | 0 | 0 | 0 | 0 | 0 | 0 | 930 |
| 190 | 3 | 3 | 3 | 1 | 0 | 0 | 0 | 0 | 0 | 0 | 4,600 |
| 220 | 3 | 3 | 3 | 3 | 2 | 0 | 0 | 0 | 0 | 0 | 110,000 |
| 260 | 3 | 3 | 0 | 1 | 0 | 0 | 0 | 0 | 0 | 0 | 380 |
| 295 | 3 | 2 | 1 | 1 | 0 | 0 | 0 | 0 | 0 | 0 | 200 |

**Supplementary Table 11.** Most probable number of aerobic chemoorganohetarotrophic microorganisms along the explored sediment-depths of SSK42/6.

| **Sediment-depth explored**  **(in cmbsf)** | **Sample dilution grade tested and the number of tubes showing positive growth response therein** | | | | | | | | | | **MPN**  **g^-1^ sediment** |
| --- | --- | --- | --- | --- | --- | --- | --- | --- | --- | --- | --- |
|  | **10^-1^** | **10^-2^** | **10^-3^** | **10^-4^** | **10^-5^** | **10^-6^** | **10^-7^** | **10^-8^** | **10^-9^** | **10^-10^** |  |
| 2 | 3 | 3 | 3 | 3 | 3 | 3 | 3 | 0 | 0 | 0 | 11,000,000 |
| 15 | 3 | 3 | 3 | 3 | 3 | 3 | 1 | 0 | 0 | 0 | 4,600,000 |
| 30 | 3 | 3 | 3 | 3 | 3 | 3 | 2 | 0 | 0 | 0 | 11,000,000 |
| 45 | 3 | 3 | 3 | 3 | 3 | 3 | 1 | 0 | 0 | 0 | 4,600,000 |
| 60 | 3 | 3 | 3 | 3 | 3 | 3 | 0 | 0 | 0 | 0 | 1,100,000 |
| 75 | 3 | 3 | 3 | 3 | 3 | 3 | 1 | 0 | 0 | 0 | 4,600,000 |
| 90 | 3 | 3 | 3 | 3 | 3 | 2 | 1 | 0 | 0 | 0 | 1,500,000 |
| 105 | 3 | 3 | 3 | 3 | 3 | 3 | 1 | 0 | 0 | 0 | 4,600,000 |
| 120 | 3 | 3 | 3 | 3 | 3 | 3 | 1 | 0 | 0 | 0 | 4,600,000 |
| 135 | 3 | 3 | 3 | 3 | 3 | 3 | 2 | 0 | 0 | 0 | 11,000,000 |
| 145 | 3 | 3 | 3 | 3 | 3 | 0 | 0 | 0 | 0 | 0 | 110,000 |
| 160 | 3 | 3 | 3 | 3 | 0 | 0 | 0 | 0 | 0 | 0 | 24,000 |
| 175 | 3 | 3 | 3 | 3 | 3 | 3 | 2 | 0 | 0 | 0 | 11,000,000 |
| 190 | 3 | 3 | 3 | 3 | 3 | 2 | 0 | 0 | 0 | 0 | 1,100,000 |
| 205 | 3 | 3 | 3 | 3 | 3 | 3 | 3 | 3 | 0 | 0 | 110,000,000 |
| 220 | 3 | 3 | 3 | 2 | 1 | 0 | 0 | 0 | 0 | 0 | 15,000 |
| 235 | 3 | 3 | 3 | 3 | 0 | 0 | 0 | 0 | 0 | 0 | 24,000 |
| 250 | 3 | 3 | 3 | 3 | 1 | 2 | 0 | 0 | 0 | 0 | 120,000 |
| 265 | 3 | 3 | 0 | 0 | 0 | 0 | 0 | 0 | 0 | 0 | 230 |
| 270 | 3 | 3 | 3 | 3 | 3 | 3 | 0 | 1 | 0 | 0 | 3,800,000 |
| 275 | 3 | 3 | 3 | 3 | 3 | 3 | 1 | 1 | 0 | 0 | 7,500,000 |

**Supplementary Table 12.** Most probable number of aerobic chemolithoautotrophic microorganisms along the explored sediment-depths of SSK42/5.

| **Sediment-depth explored**  **(in cmbsf)** | **Sample dilution grade tested and the number of tubes showing positive growth response therein** | | | | | | | | | | **MPN**  **g^-1^ sediment** |
| --- | --- | --- | --- | --- | --- | --- | --- | --- | --- | --- | --- |
|  | **10^-1^** | **10^-2^** | **10^-3^** | **10^-4^** | **10^-5^** | **10^-6^** | **10^-7^** | **10^-8^** | **10^-9^** | **10^-10^** |  |
| 0 | 2 | 0 | 1 | 0 | 0 | 0 | 0 | 0 | 0 | 0 | 14 |
| 15 | 3 | 1 | 1 | 0 | 0 | 0 | 0 | 0 | 0 | 0 | 75 |
| 45 | 3 | 3 | 0 | 1 | 0 | 0 | 0 | 0 | 0 | 0 | 380 |
| 60 | 3 | 3 | 1 | 0 | 0 | 0 | 0 | 0 | 0 | 0 | 430 |
| 90 | 3 | 3 | 2 | 1 | 0 | 0 | 0 | 0 | 0 | 0 | 1,500 |
| 120 | 3 | 3 | 2 | 0 | 0 | 0 | 0 | 0 | 0 | 0 | 930 |
| 140 | 3 | 1 | 2 | 0 | 0 | 0 | 0 | 0 | 0 | 0 | 1,200 |
| 160 | 3 | 3 | 2 | 2 | 0 | 0 | 0 | 0 | 0 | 0 | 2,100 |
| 190 | 3 | 3 | 2 | 0 | 0 | 0 | 0 | 0 | 0 | 0 | 930 |
| 220 | 3 | 3 | 1 | 3 | 0 | 0 | 0 | 0 | 0 | 0 | 1,600 |
| 260 | 3 | 2 | 1 | 0 | 0 | 0 | 0 | 0 | 0 | 0 | 150 |
| 295 | 3 | 2 | 0 | 0 | 0 | 0 | 0 | 0 | 0 | 0 | 93 |

**Supplementary Table 13.** Most probable number of aerobic chemolithoautotrophic microorganisms along the explored sediment-depths of SSK42/6.

| **Sediment-depth explored**  **(in cmbsf)** | **Sample dilution grade tested and the number of tubes showing positive growth response therein** | | | | | | | | | | **MPN**  **g^-1^ sediment** |
| --- | --- | --- | --- | --- | --- | --- | --- | --- | --- | --- | --- |
|  | **10^-1^** | **10^-2^** | **10^-3^** | **10^-4^** | **10^-5^** | **10^-6^** | **10^-7^** | **10^-8^** | **10^-9^** | **10^-10^** |  |
| 2 | 1 | 0 | 0 | 0 | 0 | 0 | 0 | 0 | 0 | 0 | 36 |
| 15 | 0 | 0 | 1 | 0 | 0 | 0 | 0 | 0 | 0 | 0 | 3 |
| 30 | 0 | 1 | 0 | 0 | 0 | 0 | 0 | 0 | 0 | 0 | 3 |
| 45 | 3 | 3 | 3 | 1 | 0 | 0 | 0 | 0 | 0 | 0 | 4,300 |
| 60 | 3 | 1 | 0 | 0 | 0 | 0 | 0 | 0 | 0 | 0 | 43 |
| 75 | 3 | 1 | 0 | 0 | 0 | 0 | 0 | 0 | 0 | 0 | 43 |
| 90 | 3 | 3 | 3 | 3 | 2 | 3 | 0 | 0 | 0 | 0 | 290,000 |
| 105 | 3 | 3 | 3 | 2 | 1 | 1 | 0 | 0 | 0 | 0 | 20,000 |
| 120 | 3 | 3 | 2 | 0 | 0 | 0 | 0 | 0 | 0 | 0 | 930 |
| 135 | 3 | 2 | 3 | 0 | 1 | 1 | 0 | 0 | 0 | 0 | 6,100 |
| 145 | 3 | 3 | 3 | 3 | 0 | 0 | 0 | 0 | 0 | 0 | 11,000 |
| 160 | 3 | 3 | 3 | 3 | 3 | 2 | 0 | 1 | 0 | 0 | 1,400,000 |
| 175 | 3 | 3 | 3 | 2 | 3 | 1 | 2 | 1 | 0 | 0 | 1,500,000 |
| 190 | 3 | 3 | 3 | 3 | 3 | 3 | 0 | 0 | 0 | 0 | 1,100,000 |
| 205 | 3 | 3 | 3 | 3 | 3 | 1 | 0 | 0 | 0 | 0 | 460,000 |
| 220 | 3 | 3 | 3 | 3 | 3 | 3 | 3 | 1 | 0 | 0 | 46,000,000 |
| 235 | 3 | 3 | 3 | 3 | 3 | 3 | 1 | 0 | 0 | 0 | 4,600,000 |
| 250 | 3 | 3 | 3 | 3 | 3 | 3 | 1 | 0 | 0 | 0 | 4,600,000 |
| 265 | 3 | 3 | 3 | 3 | 3 | 3 | 1 | 0 | 0 | 0 | 4,600,000 |
| 270 | 3 | 3 | 3 | 3 | 3 | 3 | 2 | 0 | 0 | 0 | 11,000,000 |
| 275 | 3 | 3 | 3 | 3 | 3 | 3 | 2 | 0 | 0 | 0 | 11,000,000 |

**Supplementary Table 31.** Growth behavior of *Halothiobacillus* sp. SB14A in co-cultures with or without perchlorate-respiring bacterial consortia (PRBC), in a consensus medium supplemented with or without perchlorate and containing energy, electron and carbon sources suitable for both *Halothiobacillus* and perchlorate-respirers.

|  | **I. Growth in consensus medium supplemented with perchlorate** | | | | |
| --- | --- | --- | --- | --- | --- |
|  | SB14A culture not accompanied by PRBC | SB14A in co-cultures with PRBC | | | |
|  |  | PRBC source: 120 cmbsf of SSK42/5 | PRBC source: 260 cmbsf of SSK42/5 | PRBC source: 135 cmbsf of SSK42/6 | PRBC source: 275 cmbsf of SSK42/6 |
|  | **Ia. Anaerobic incubation** | | | | |
| *Halothiobacillus* CFU mL^-1^ medium at 0 h of incubation | 8.7*10^4^ | 8*10^4^ | 2*10^4^ | 6*10^4^ | 1.5*10^4^ |
| *Halothiobacillus* CFU mL^-1^ medium after 30 days’ incubation | 2.6*10^3^ | 5.3*10^5^ | 2.1*10^4^ | 1*10^4^ | 3.5*10^4^ |
| *Halothiobacillus* CFU mL^-1^ medium after 60 days’ incubation | 3.5*10^2^ | 1.8*10^5^ | 1.3*10^5^ | 7*10^4^ | 3.3*10^5^ |
|  | **Ib. Aerobic incubation** | | | | |
| *Halothiobacillus* CFU mL^-1^ medium at 0 h of incubation | 6.5*10^5^ | 3*10^5^ | 7.7*10^5^ | 5.7*10^5^ | 6.7*10^5^ |
| *Halothiobacillus* CFU mL^-1^ medium after 2 days’ incubation | 2.9*10^8^ | 3.7*10^8^ | 5.9*10^8^ | 6.5*10^8^ | 8.3*10^8^ |
|  | **II. Growth in consensus medium having no perchlorate** | | | | |
|  | SB14A culture not accompanied by PRBC | SB14A in co-cultures with PRBC | | | |
|  |  | PRBC source: 120 cmbsf of SSK42/5 | PRBC source: 260 cmbsf of SSK42/5 | PRBC source: 135 cmbsf of SSK42/6 | PRBC source: 275 cmbsf of SSK42/6 |
|  | **IIa. Anaerobic incubation** | | | | |
| *Halothiobacillus* CFU mL^-1^ medium at 0 h of incubation | 7.9*10^4^ | 6.7*10^4^ | 7.1*10^4^ | 6.9*10^4^ | 7.5*10^4^ |
| *Halothiobacillus* CFU mL^-1^ medium after 60 days’ incubation | 3.1*10^2^ | 2.6*10^2^ | 2.5*10^2^ | 2.1*10^2^ | 3*10^2^ |

**Supplementary Table 32.** (Per)chlorate reductase genes identified within the contigs assembled from the metatranscriptomic sequence data obtained from the 275 cmbsf sediment-sample of SSK42/6.

| **Serial**  **No.** | **Query name** | **Seed eggNOG ortholog** | **Organismal source of the closest hit** | **Seed ortholog evalue** | **Seed ortholog score** | **KEGG Orthology Numbers** | **COG cat** | **eggNOG annotation** |
| --- | --- | --- | --- | --- | --- | --- | --- | --- |
| 1 | ALNBEFHC_07263 | 351348.Maqu_3086 | *Marinobacter hydrocarbonoclasticus* VT8 | 0 | 2537.7 | K17050, K00370 | C | Chlorate reductase subunit alpha, nitrate reductase subunit alpha |
| 2 | ALNBEFHC_07262 | 290398.Csal_1331 | *Chromohalobacter salexigens* DSM 3043 | 3.1E-261 | 906 | K17051, K00371 | C | Chlorate reductase subunit beta, nitrate reductase subunit beta |

**Supplementary Methods**

**Determining Most Probable Number (MPN) of aerobic, chemoorganoheterotrophs and chemolithoautotrophs.** MPN of viable cells of aerobic, chemoorganoheterotrophs and sulfur-chemolithoautotrophs in the individual sediment-samples were calculated using ten-fold dilution series (and three tubes per dilution) of LB or ASWT slurry-cultures respectively^1^. For each community, 1 g sediment-sample was dissolved in 10 mL sterilized 0.9% (w/v) NaCl and vortexed for 2-3 min. The sediment slurry was placed on a 15°C rotary incubator (150 rpm) for 2 h. Subsequently, the slurry was kept undisturbed for 1 h; after all the visible sediment particles had settled down, 1mL of this 10^-1^ dilution-grade (0.1 g sediment mL^-1^) was transferred to another tube containing 9 mL 0.9% (w/v) NaCl, thereby generating the 10^-2^ dilution-grade (0.01 g sediment mL^-1^). For seven further iterations, inoculum from the last dilution-grade was transferred to fresh tubes containing 9 mL 0.9% (w/v) NaCl, so as to generate a dilution series up to 10^-9^. Finally, from each dilution-grade, three 0.5 mL aliquots of particle-free suspension were transferred to a triplicate set of MPN-test tubes containing 4.5 mL of the sterilized medium of choice. In doing so, the 10^-1^ to 10^-9^ dilution-grades created in 0.9% (w/v) NaCl gave rise to nine sets of MPN-test tube triplicates that represented the inoculum-dilution grades 10^-2^ to 10^-10^; so MPN-test tubes representing the 10^-1^ inoculum-dilution grade was generated separately by adding 0.5 g sediment-sample directly to 5 mL sterilized medium. All MPN-test tubes were mixed thoroughly, and incubated aerobically at 15°C, for 72 h and 21 days according as they contained LB or ASWT. LB culture tubes which showed turbidity against the blank were regarded as positive for chemoorganoheterotrophic growth while ASWT culture tubes that exhibited acidification (by changing the color of phenol red indicator present in the medium) were regarded as positive for growth of chemolithoautotrophs. Growth responses observed in the MPN-tube series were tallied with a standard MPN Table^1^ (https://www.fda.gov/food/foodscienceresearch/laboratorymethods/ucm109656.htm) to get the MPN of the metabolic type g^-1^ sediment.

**Composition of the media used in this study.** 1 L Luria-Bertani (LB) broth contained 10 g tryptone, 5 g yeast extract and 5 g NaCl. 1 L Reasoner’s 2A (R2A) broth contained 0.5 g proteose peptone, 0.5 g casamino acids, 0.5 g yeast extract, 0.5 g dextrose, 0.5 g soluble starch, 0.3 g K_2_HPO_4_, 0.05 g MgSO_4_.7H_2_O, 0.3 g sodium pyruvate.

MSTY medium contained modified basal and mineral salts (MS) solution supplemented with 20 mM Na_2_S_2_O_3_.5H_2_O and 500 mg yeast extract L^-1^. MS contained the following L^-1^ distilled water: 1 g NH_4_Cl, 4 g K_2_HPO_4_, 1.5 g KH_2_PO_4_, 0.5 g MgSO_4_.7H_2_O and 5.0 mL trace metals solution^2^. 1 L trace element solution (pH 6.0) contained 50 g EDTA, 22 g ZnSO_4_.7H_2_O, 5.06 g MnCl_2_, 4.99 g FeSO_4_, 1.1 g (NH_4_)_6_ MoO_26_.4H_2_O, 1.57 g CuSO_4_ and 1.61 g CoCl_2_.6H_2_O.

ASWT medium contained artificial sea water (ASW) supplemented with 20 mM Na_2_S_2_O_3_.5H_2_O (added separately after filter sterilization)^3^. ASW contained the following L^-1^ distilled water: 25.1 g NaCl, 1 g (NH_4_)_2_SO_4_, 1.5 g MgSO_4_, 7H_2_O, 0.3 g CaCl_2_.2H_2_O, 0.2 g NaHCO_3_, 2.4 g Tris, 1 mL trace element solution and 0.5 g K_2_HPO_4_ (added after autoclaving separately). 1 L trace element solution (pH 6.0) contained 50 g EDTA, 22 g ZnSO_4_.7H_2_O, 5.06 g MnCl_2_, 4.99 g FeSO_4_, 1.1 g (NH_4_)_6_ MoO_26_.4H_2_O, 1.57 g CuSO_4_ and 1.61 g CoCl_2_.6H_2_O.

1 L perchlorate-supplemented, basal bicarbonate-buffered (PBBB) medium contained 0.53 g Na_2_HPO_4_.2H_2_O, 0.41 g K_2_HPO_4,_ 0.3 g NH_4_Cl, 0.11 g CaCl_2_.2H_2_O, 0.10 g MgCl_2_.6H_2_O, 0.3 g NaCl, 4 g NaHCO_3_, 0.48 g Na_2_S, 50 mg yeast extract, 1.36 g CH_3_COONa, 2 mL trace metal solution, 20 mM CH_3_OH and 20 mM NaClO_4_, and was supplemented with 0.5 g L^-1^ sodium thioglycolate and 0.1 mg L^-1^ resazurin. Here, 1 L trace element solution (pH 6.0) contained 50 g EDTA, 22 g ZnSO_4_.7H_2_O, 5.06 g MnCl_2_, 4.99 g FeSO_4_, 1.1 g (NH_4_)_6_ MoO_26._4H_2_O, 1.57 g CuSO_4_ and 1.61 g CoCl_2._6H_2_O.

1 L PBBB-ASWT consensus medium contained 5 g NaCl, 1.5 g MgSO_4._ 7H_2_O, 0.2 g CaCl_2_.2H_2_O, 1 g (NH_4_)_2_SO_4_, 1 g CH_3_COONa, 2 g Tris, 0.53 g Na_2_HPO_4_.2H_2_O, 20 mg yeast extract, 2.5 g Na_2_S_2_O_3_, 20 mM NaClO_4_, 1 g NaHCO_3_, 0.45 g K_2_HPO_4_ and 2 mL trace metal solution (pH 6.0), every liter of which contained 50 g EDTA, 22 g ZnSO_4_.7H_2_O, 5.06 g MnCl_2_, 4.99 g FeSO_4_, 1.1 g (NH_4_)_6_ MoO_26._4H_2_O, 1.57 g CuSO_4_, 1.61 g CoCl_2._6H_2_O.

**Metagenome (total community DNA) sequencing.** Quality of metagenomic DNA samples was checked by electrophoresis and considered to be of high quality when no degradation signs were apparent. DNA quantity was determined using Qubit dsDNA HS Assay Kit (Thermo Fisher Scientific). 1 μg DNA from each sediment-sample was taken for deep shotgun sequencing by the Ion Proton platform using 200 bp read chemistry on a PI V2 Chip.

Libraries to be used for sequencing were constructed using the Ion Plus Fragment Library Kit (Thermo Fisher Scientific) and following the manufacturer’s Ion Plus gDNA and Amplicon Library Preparation User Guide. The Proton library was generated using 1 µg of genomic DNA which was fragmented to approximately 200 base pairs by the Covaris S2 system (Covaris, Inc., USA) and purified with 1.8x Agencourt Ampure XP Beads (Beckman Coulter, USA). Fragmentation was followed by end-repair, blunt-end ligation of the Ion Xpress Barcode and Ion P1 adaptors, and nick translation.

Post-ligation, size selection was done using E-Gel Size-Select 2% Agarose gels (Thermo Fisher Scientific) with the target size of 300 bp. Final PCR was performed using platinum PCR SuperMix High Fidelity and Library Amplification Primer Mix (Thermo Fisher Scientific), for 5 cycles of amplification. The resulting library was purified using AMPure XP reagent (1.2x; Beckman Coulter) and the concentration determined with Qubit dsDNA HS Assay Kit (Thermo Fisher Scientific); size distribution was done with Agilent 2100 Bioanalyzer high-sensitivity DNA kit (Agilent Technologies). Libraries were pooled in equimolar concentrations and used for template preparation.

Library templates were prepared for sequencing using OneTouch 2 protocols and reagents (Thermo Fisher Scientific). Library fragments were clonally amplified onto ion sphere particles (ISPs) through emulsion PCR and then enriched for template-positive ISPs. Proton emulsion PCR reactions utilized the Ion PI Template OT2 200 Kit v3 (Thermo Fisher Scientific). Following recovery, enrichment was completed by selectively binding the ISPs containing amplified library fragments to streptavidin coated magnetic beads, removing empty ISPs through washing steps, and denaturing the library strands to allow for collection of the template-positive ISPs. For all reactions, these steps were accomplished using the ES module of the Ion OneTouch 2. The selected ISPs were loaded on PI V2 Chip and sequencing was performed with the Ion PI 200 Sequencing Kit (Thermo Fisher Scientific) using the 500-flow (125 cycle) run format.

**Whole genome sequencing of the bacterial isolates.** Quality of the genomic DNA samples extracted from the 6 bacterial isolates were checked by electrophoresis on 1% (w/v) agarose gel and considered to be of good quality if degradation was not observed. Qubit dsDNA HS Assay Kit (Thermo Fisher Scientific) was then used to determine DNA quantity. 100 ng DNA from each isolate was taken for deep shotgun sequencing on an Ion S5 next-generation sequencing platform (Thermo Fisher Scientific) using 400 bp read chemistry on a 530 or 520 Chip.

Libraries were constructed by the Ion Xpress Plus Fragment library kit (Thermo Fisher Scientific) using 100 ng genomic DNA from each isolate. In this procedure genomic DNA samples were fragmented using Ion Shear Plus Reagents (Thermo Fisher Scientific) which enzymatically fragment DNA. The fragmented libraries were then purified by 1.8X Agencourt Ampure XP Beads (Beckman Coulter). These purified libraries were then subjected to barcode-adapter ligation and nick repair. Adapter-ligated and nick-repaired libraries were purified again by 1X Agencourt Ampure XP Beads (Beckman Coulter).

Size selection of the libraries was done using E-Gel Size-Select 2% Agarose gels (Thermo Fisher Scientific) with 480 bp target size. Final PCR was performed using platinum SuperMix High Fidelity PCR system and Library Amplification Primer Mix (both from Thermo Fisher Scientific), for 8 cycles of amplification. The resulting libraries were purified using 1x Agencourt AMPure XP reagent (Beckman Coulter). Concentrations of the purified libraries were determined with Qubit dsDNA HS Assay Kit (Thermo Fisher Scientific). Libraries were then pooled in equimolar concentrations and used for template preparation.

The library template to be used for sequencing was prepared using Ion OneTouch 2 reagents (Thermo Fisher Scientific). Library fragments were clonally amplified onto ion sphere particles (ISPs) through emulsion PCR and then enriched for template-positive ISPs. Following emulsion PCR, enrichment was completed by selectively binding the ISPs containing amplified library fragments to streptavidin coated magnetic beads, removing empty ISPs through washing steps, and denaturing the library strands to allow for collection of the template-positive ISPs using the Ion OneTouch ES instrument (Thermo Fisher Scientific). The selected ISPs were loaded on a 530 or 520 Chip and sequencing was performed with the Ion S5 Sequencing Kit (Thermo Fisher Scientific) using the 850-flow run format.

**Metatranscriptome sequencing.** Metatranscriptome was extracted from the 275 cmbsf sediment-sample of SSK42/6 using the RNA PowerSoil Total RNA Isolation Kit (MoBio), as per manufacturer’s protocol. Nanogram level total RNA was obtained after pooling up the products of 15 individual preparatory reactions, each carried out using 2 g sediment-sample. Individual total RNA preparations were all subjected to DNase digestion by RNase free DNase I (Thermo Fisher Scientific) and purified using RNeasy MinElute Cleanup Kit (Qiagen, Germany); their concentrations were measured using Quant-iT RiboGreen RNA Assay Kit (Thermo Fisher Scientific). To assess the integrity of total RNA (RIN) in these preparations, their aliquots were run on a TapeStation RNA ScreenTape electrophoretic system (Agilent Technologies) and only high-quality preparations having RIN value >7.0 were added to the RNA pool to be used for library construction. The mRNA species thus pooled up were selectively converted into a library of template molecules using TruSeq Stranded mRNA and Total RNA kit (Illumina Inc., USA). Depletion of rRNAs was carried out using the Ribo-Zero Gold system (Illumina), which is an integral part of the kit used for preparing the library. The mRNAs were fragmented into small pieces using divalent cations under elevated temperature. The cleaved RNA fragments were copied into first strand cDNAs using reverse transcriptase and random primers. This was followed by second strand cDNA synthesis using DNA Polymerase I and RNase H. cDNA fragments were then subjected to end-repair, addition of single ‘A’ bases, adaptor ligation, purification and enrichment with PCR to create the final library, which was sequenced using a HiSeq4000 (Illumina).

**Supplementary References**

1.	Sutton, S. The most probable number method and its uses in enumeration, qualification, and validation. *Microbiol*. *Top*. *J*. *Validation Tech*. **16,** 35-38 (2010).

2.	Ghosh, W. & Roy, P. *Mesorhizobium* *thiogangeticum* sp. nov., a novel sulfur-oxidizing chemolithoautotroph from rhizosphere soil of an Indian tropical leguminous plant. *Int*. *J*. *Syst*. *Evol*. *Microbiol*. **56,** 91-97 (2006).

3.	Alam, M., Pyne, P., Mazumdar, A., Peketi, A. & Ghosh, W. Kinetic enrichment of ^34^S during proteobacterial thiosulfate oxidation and the conserved role of SoxB in S-S bond breaking. *Appl*. *Environ*. *Microbiol*. **79,** 4455-4464 (2013).
